# Place cell activity and behaviour during task acquisition and extinction learning predict renewal outcome

**DOI:** 10.1101/2025.07.07.663428

**Authors:** Laura Dolón-Vera, José Donoso, Sen Cheng, Denise Manahan-Vaughan

**Affiliations:** Department of Neurophysiology, Medical Faculty, Ruhr University Bochum, Germany; Institute of Neural Computation, Faculty of Computer Science, Ruhr University Bochum, Germany

**Keywords:** hippocampus, extinction learning, place cells, spatial learning, procedural learning, rodent

## Abstract

Extinction learning (EL) and the subsequent renewal of learned behaviours are crucial for adaptive responding, yet the underlying neural mechanisms that differentiate successful renewal from its absence remain unclear. Here, we explored the behavioral and neurophysiological basis of spatial appetitive EL, as well as renewal failure and renewal success. We recorded place cell activity from hippocampal area CA1 in male rats that performed a context-dependent spatial appetitive learning task in a T-maze (rewarded context A), followed by EL (unrewarded context B), and subsequent renewal testing (unrewarded context A). Half of the animals exhibited significant renewal (“renewers”) and the other half failed to renew (“non-renewers”). Our findings reveal fundamental differences in learning strategies between groups revealed by differences in both spatial behavior and in place cell activity during both initial acquisition and subsequent EL. Specifically, renewers exhibited a context-based EL strategy, whereas non-renewers followed a goal-directed strategy. The spatial distribution of hippocampal place cell activity differed significantly between groups, indicating the hippocampus’ role in the learning processes. Furthermore, renewers exhibited a greater extent of global remapping of place cells compared to non-renewers, consistent with predictions by our computational modeling. This suggests that global remapping serves as a key neural mechanism underlying effective renewal, allowing for the segregation of distinct memories and preventing the generalization of extinction. Our results highlight how hippocampal place cell dynamics during acquisition and EL predict later behavioral renewal outcomes, providing critical insights into the neural basis of memory updating and contextual control over learned behaviors.

**Significance Statement:** This study reveals that individual differences in associative learning strategies determine the outcome of memory renewal/re-activation. After first learning, and then extinguishing, a spatial learning task in two different contexts, some animals renewed their behavior when returned to the acquisition context (“renewers”), while others did not (“non-renewers”). Intriguingly, hippocampal place cell activity during acquisition and extinction learning predicted the subsequent renewal efficacy in these distinct animal groups. In particular, renewers exhibited more global remapping than non-renewers, a neural mechanism that segregates memories from different contexts and prevents over-generalization of extinction. These findings provide critical insights into how the hippocampus supports context-dependent memory acquisition and updating, providing new insights into the physiological basis of individual differences in behavioral flexibility.

## Introduction

During extinction learning (EL) an individual learns that previously learned associations are no longer behaviorally relevant. Forms of EL that are aversive, and/or fear-related, involve activation of brain regions and circuitry that support stimulus-response based learning (1, 2), which typically falls into the category of implicit learning (3). In rodents, a change of environmental context greatly expedites spatial appetitive EL (4), but also renders it vulnerable to renewal effects when animals are returned to the original context in which the appetitive behavior was learned (4). In therapeutic settings, behavioral renewal (i.e. reactivation) can be an undesirable outcome, especially if the goal is to sustain the continuation of extinguished behavior of, for example, poor dietary habits. However, in spatial appetitive forms of EL, behavioral renewal can be advantageous.

The hippocampus supports EL of aversive experience (1). This presumably relates to the engagement of the hippocampus in conditioned, or context-dependent associative learning of aversive experience (5). However, the hippocampus is also potently engaged by spatial appetitive EL (6). This is explained by the requirement of the hippocampus for the acquisition and encoding of spatial experience (7), but also by its modulation by reward structures such as the ventral tegmental area (8, 9). To date, however, real-time electrophysiological recordings from the hippocampus during spatial appetitive context-dependent EL, have not been conducted. Thus, its precise functional contribution to this process is unclear. Furthermore, although new inhibitory learning is believed to be a component of EL of aversive experience (3), it is less clear whether it is an intrinsic component of hippocampus-dependent EL.

In this study we conducted single-unit recordings from place cells in rat hippocampus (dorsal CA1 region) during context-dependent spatial appetitive EL and subsequent renewal. Our goals were: **1**. To explore the neurophysiological and behavioral fundaments of successful and failed context-dependent renewal. **2**. To examine how the hippocampus becomes engaged in spatial appetitive EL and renewal. **3**. To leverage place cell recordings as a means to interpret the extent to which information updating and/or new learning support spatial appetitive EL. By this means, we tested the hypothesis that the degree and precise nature of hippocampal activation during spatial appetitive EL potently influences renewal efficacy.

## Results

### Some animals show renewal, others do not

Rats (n = 10) participated in an ABA paradigm in which task acquisition (context A), EL (context B) and subsequent behavioral renewal (triggered by exposure to the original A context) were assessed (Fig. 1A). Contexts differed by floor patterns of the T-maze, subtle odor cues at the ends of the arms, and distal wall cues (10). A food reward was placed at the end of the right arm of the T-maze during acquisition. Animals performed 20 trials/day (4 sessions, 5 trials/day) for 3 days, with reward probability gradually reduced from 100% (first 10 trials on day 1) to 30% (last 10 trials on day 3). All animals reached >80% correct arm choices by the end of acquisition in context A (Fig. 1B, Sessions 1-12; SI Appendix Table S1). During EL on day 4 in context B (unrewarded), arm choice performance returned to chance levels (Fig. 1B, S13-15). When subsequently tested for renewal in context A (unrewarded), some animals (*n* = 5, “renewers”) maintained >80% correct choices, whereas others (n=5, “non-renewers”) did not (chance level) (Fig. 1C, S16; SI Appendix Table S1). There were no group differences in alternative responses during acquisition, but a significant difference emerged during the renewal phase (SI Appendix Fig. S1A, Table S1).

**Figure 1.**
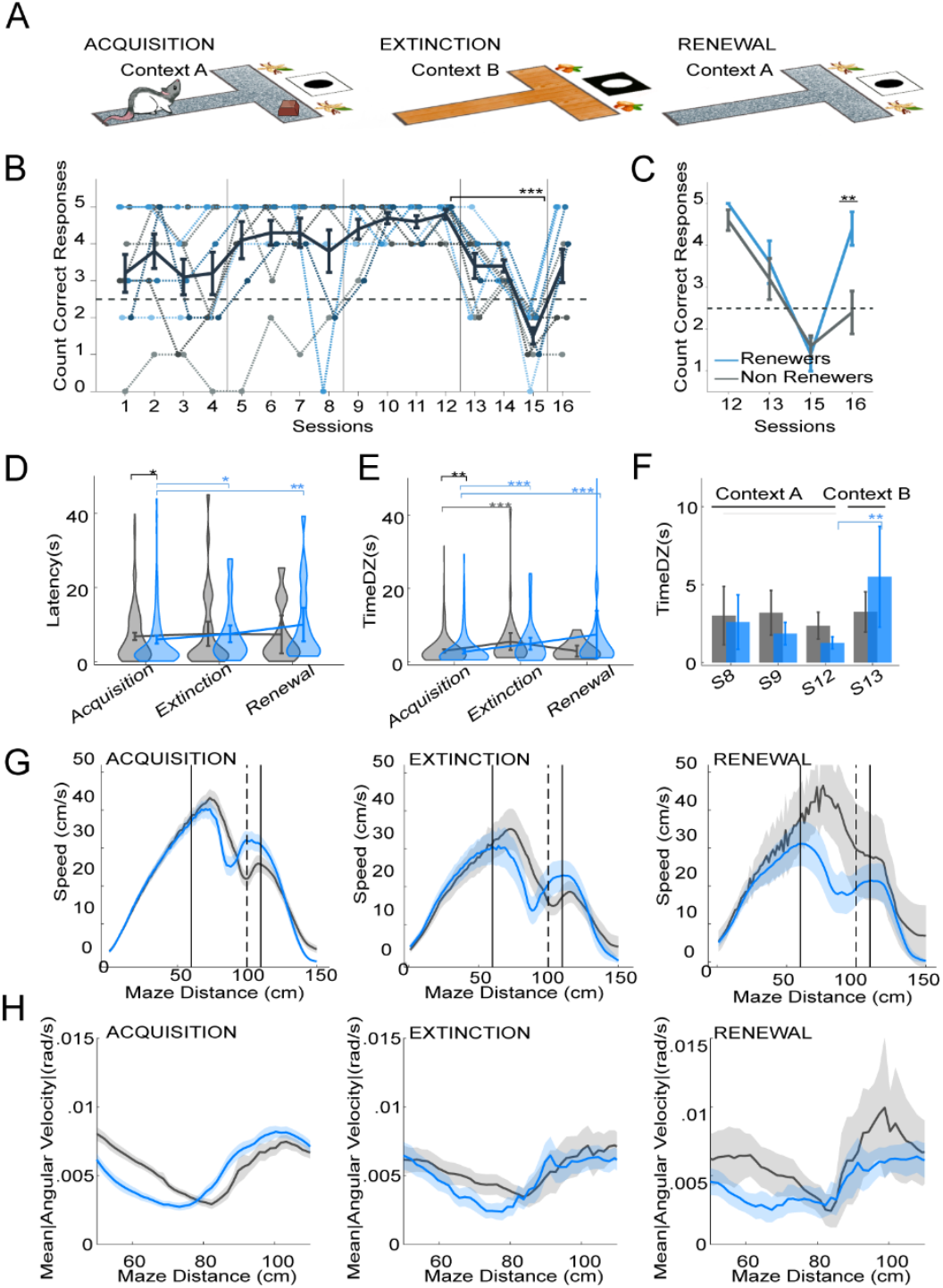
Appetitive spatial ABA renewal paradigm revealed two distinct learning strategies. **(A)** Rats were trained for 3 days in a rewarded T-maze (context A, acquisition), underwent extinction learning (EL) in an unrewarded, distinct, context B, and were tested for renewal in context A without reward. **(B)** Correct arm choices across sessions (5 trials/session). Dashed lines represent individual animals. Grey dashed line represents chance (2.5). n=10 animals acquired the task (Sessions 1–12) and extinguished (Sessions 13–15); only half displayed renewal (Session 16). **(C)** Correct choices split by group: renewers (blue) and non-renewers (grey). **(D–E)** Violin plots showing latencies. **(D)** and decision zone (DZ) dwell times **(E)** during correct responses. **(F)** Bar plot of DZ times across key sessions (last Day 2 [S8], first Day 3 [S9], last Day 3 [S12], first Day 4 [S13]). **(G–H)** Line plots of maze speed **(G)** and angular velocity **(H)** across phases (**see also SI Appendix, Fig. S1**). Maze divided into: S (0–20), corridor (21–60), junction (61–100), intersection (101–110), and right arm (111–150). Error bars show mean ± SEM (B–C) or mean ± 95% confidence interval (CI) (D–H). Asterisks indicate significance (*p<0.05): blue, within renewers across phases; grey, within non-renewers; black, between groups.

### Renewers and non-renewers exhibit different behaviour during acquisition and EL

We next examined the animals’ behavior to study the learning strategies adopted by renewers and non-renewers. Latencies (time elapsed from the opening of the door till animals leave the starting zone, S), time spent in decision zones (junction -J- and intersection -I-, DZ) prior to arm choice, and dwell times in DZ after correct responses were analysed (Fig. 1E and SI Appendix Fig. S1B-F). During acquisition, renewers exited S more quickly and spent less time in DZ prior to arm choices than non-renewers (Fig. 1D-E; *SI Appendix*, Table S1), suggesting faster task engagement and earlier decision-making. These differences were not observed during EL or renewal. Interestingly, non-renewers spent more time in DZ *after* correct arm choices in unrewarded trials, and less time after correct arm choices that were rewarded, suggesting a higher sensitivity of non-renewers to reward history (*SI Appendix* Fig. S1C, Table S1).

Later, renewers showed increased latencies and dwell times from acquisition to EL and renewal, whereas non-renewers did not (Fig. 1D-E; *SI Appendix* Table S1). During early EL (S13), only renewers spent significantly more time in DZ compared to the last acquisition session (S12), suggesting their perception of the context change (Fig. 1F; *SI Appendix* Table S1). No differences were found between consecutive acquisition sessions (S8–9), indicating that context-specific learning may have occurred in renewers (Fig. 1F). Maze speed profiles further distinguished the two groups: renewers showed a progressive slowing across learning phases, while non-renewers slowed during EL and returned to acquisition-level speeds during renewal, despite the absence of renewal behavior (Fig. 1G; SI Appendix Fig. S1D, Table S1). During acquisition, renewers decelerated earlier in J while non-renewers decelerated at I (Fig. 1G; *SI Appendix* Fig. S1E-F). These patterns persisted during EL but converged during renewal. Angular velocity analyses revealed that renewers exhibited greater directional variability earlier e.g. at trial initiation and DZ, consistent with thorough exploration (Fig. 1H; *SI Appendix*, Fig. S1G-H, Table S1). These differences suggest that different learning strategies had been adopted by the two groups, as early as during acquisition. Moreover, renewers appeared more sensitive to context changes and more flexible in their navigation, while non-renewers were more strongly driven by recent reward history and showed less behavioral adaptation.

### Distinct spatial properties emerge that differentiatie renewers from non-renewers across ABA paradigm

Next, we assessed if place cell (PC) firing behavior (*SI Appendix* Fig. S2A,B) could provide insights as to the neurophysiological basis of these behavioral differences. Following the assessment of spatial properties of the firing rate maps and quantification of spatial information content of single units recorded from dorsal CA1, we classified 379 PCs in non-renewers and 305 in renewers (*SI Appendix* Fig. S2C-F). The firing activity was projected onto a linearized position (*SI Appendix*, Fig. S2B), and one-dimensional firing rate maps were generated for each trial (Fig. 2A).

**Figure 2.**
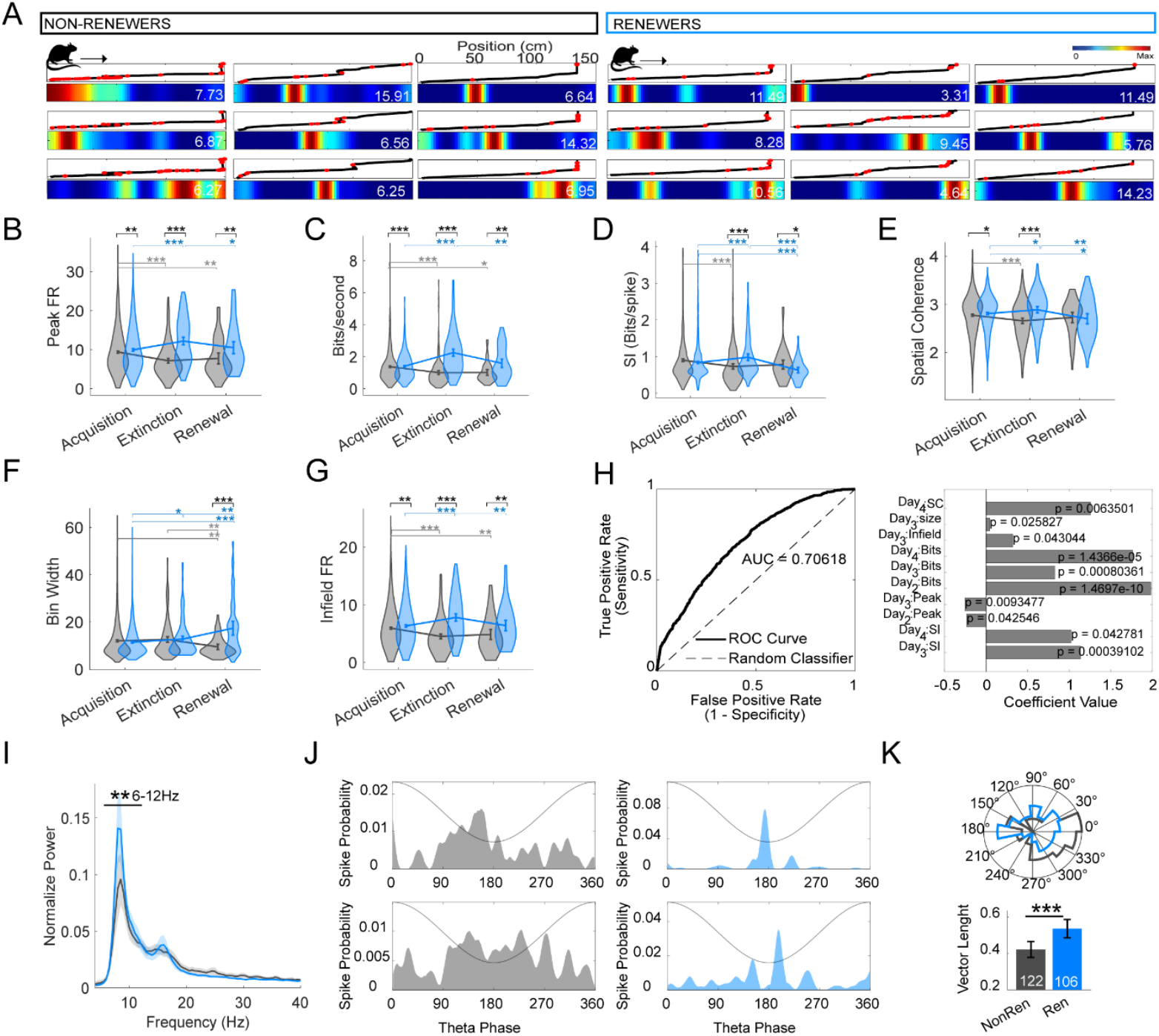
Distinct place cell coding and theta coupling underlie renewal vs. non-renewal. **A)** Examples of firing rate maps from non-renewers (left) and renewers (right). Each column shows a different trial; rows depict individual place cells recorded simultaneously. Black traces represent the linearized trajectory, red ticks mark spikes, and color scale shows normalized firing rates (white numbers indicate peak). **(B–G)** Violin plots of place cell features across learning phases: peak firing rates **(B)**, bits/s **(C)**, spatial information **(D)**, spatial coherence **(E)**, field size (width in bins, **F**), and infield firing rates (**G**). **(H)** Logistic GLM predicting renewal expression. Left, ROC curve (AUC = 0.706). Right, regression coefficients for significant predictors. **(I)** Normalised power spectra from all correct trials. **(J)** Theta phase locking for two example neurons from non-renewers (left) and renewers (right); schematic theta cycle shown in grey. **(K)** Top: polar histogram of preferred theta phases for significantly phase-locked cells. Bottom: strength of phase locking (mean resultant length). White bars indicate the number of significantly modulated cells. Error bars represent mean ± CI. Asterisks indicate significance (*p<0.05): blue, within renewers across phases; grey, within non-renewers; black, between groups.

Renewers consistently exhibited higher peak firing rates, bits/s and average firing rates than non-renewers across all test conditions (Fig. 2B-C; SI Appendix, Table S3-5), suggesting stronger spatial encoding of the T-maze contexts (11). Notably, renewers showed increased peak firing during EL compared to acquisition, whereas non-renewers displayed the opposite pattern. We further quantified spatial features of PC activity (Fig. 2D-G; SI Appendix Table S3-5) to evaluate their spatial tuning and their goal-directed activity (12, 13). During acquisition, both groups showed similar spatial information (Fig. 2D) and field size (Fig. 2F), but renewers exhibited higher spatial coherence (Fig. 2E), infield firing (Fig. 2G), and Out/In ratios (SI Appendix, Table S5), suggesting more spatially tuned representations. Across testing days, renewers showed a progressive increase in spatial information content, while non-renewers exhibited a decline along with increased Out-field/In-field firing ratios (*SI Appendix* Table S3-5). Additionally, renewers also exhibited higher spatial information than non-renewers on Day 3 of acquisition, suggesting greater reliance on spatial encoding during learning. During EL, PCs representations in the two groups diverged even more clearly. In comparison with acquisition, Non-renewers showed reduced infield firing and spatial information, suggesting a loss of goal-directed coding and spatial tuning; in contrast, renewers exhibited enhanced spatial tuning, with higher information scores and higher infield activity (Fig. 2D-G; *SI Appendix* Table S3-5). This suggests that context-specific spatial representations supported EL in renewers.

During renewal, peak and infield firing rates in renewers remained stable compared to acquisition, whereas average firing rates increased, and place fields became larger and less spatially precise, with reduced spatial information and coherence (Fig. 2A-G; *SI Appendix* Table S3-5). This suggests that representational updating occurred due to the reward absence. In contrast, PCs in non-renewers showed reduced size and infield firing rates, increased out/in-field ratios and stable spatial information and coherence. These findings suggest that PCs in renewers remained dynamically responsive across phases, with increased spatial tuning during context changes, whereas in non-renewers spatial encoding was rigid, context-invariant, and progressively degraded.

We subsequently wondered whether the spatial properties of PCs could predict renewal outcome. A logistic regression model (logistic GLM) was applied using renewal expression (‘yes’/’no’) as the outcome variable, whereas place field features across acquisition and EL phases were used as predictors (Fig. 2H). The model showed a good fit (AUC = 0.706; Fig. 2H left), indicating moderate predictive power. Key predictors included spatial information, bits per second, and infield firing rates during Days 2 and 3 of acquisition, as well as during EL, which were all elevated in renewers (Fig. 2H right). Finally, we also found that renewers exhibited a higher theta power and organisation of firing activity within theta frequency (6-12Hz) during correct trials, compared to non-renewers (Fig. 2I-K; *SI Appendix* Table S4), indicating a greater engagement of coordinated hippocampal processing during the paradigm. These results indicate that context-sensitive spatial tuning and coordinated theta activity underlies effective renewal, whereas rigid, degraded PC coding characterises renewal failure.

### Renewers exhibit a higher extent of place field remapping across contexts

The spatial firing of PCs highlights locations of behavioral relevance, such as cues or reward sites (14), and also serves as a representation of the environment (15). We hypothesised that if animals perceived context B as being distinct from context A, PCs would show changing spatial firing patterns across phases. Conversely, similar representations during acquisition and renewal would indicate context-specific reinstatement. We tested this by comparing spike distributions, place field shifts and decoding across conditions (Fig. 3). First, we analysed spike distribution (Fig. 3A). During acquisition, both groups showed spike clustering in S, DZ, and the target arm. Nonetheless, group-specific patterns emerged: non-renewers exhibited a higher spike density in S, while renewers concentrated spikes in the decision zone, suggesting reliance on different navigation strategies (Fig.3A left; *SI Appendix* Table S6). During EL, reward-related activity decreased in both groups. Yet, only renewers exhibited a global redistribution of spikes, while non-renewers retained their previous spike pattern. During renewal, non-renewers returned to their acquisition-like firing patterns, whereas renewers maintained the distributions seen during EL. Rate maps confirmed these patterns (SI Appendix Fig. S3A Table S7): PCs of renewers showed significant rate remapping between acquisition and EL phases, while non-renewers exhibited invariant spatial firing profiles throughout the acquisition and EL phases. During renewal, renewers exhibited a further increase of spiking activity in DZ, suggesting re-engagement with spatial cues and potential reinstatement of context A representations.

**Figure 3.**
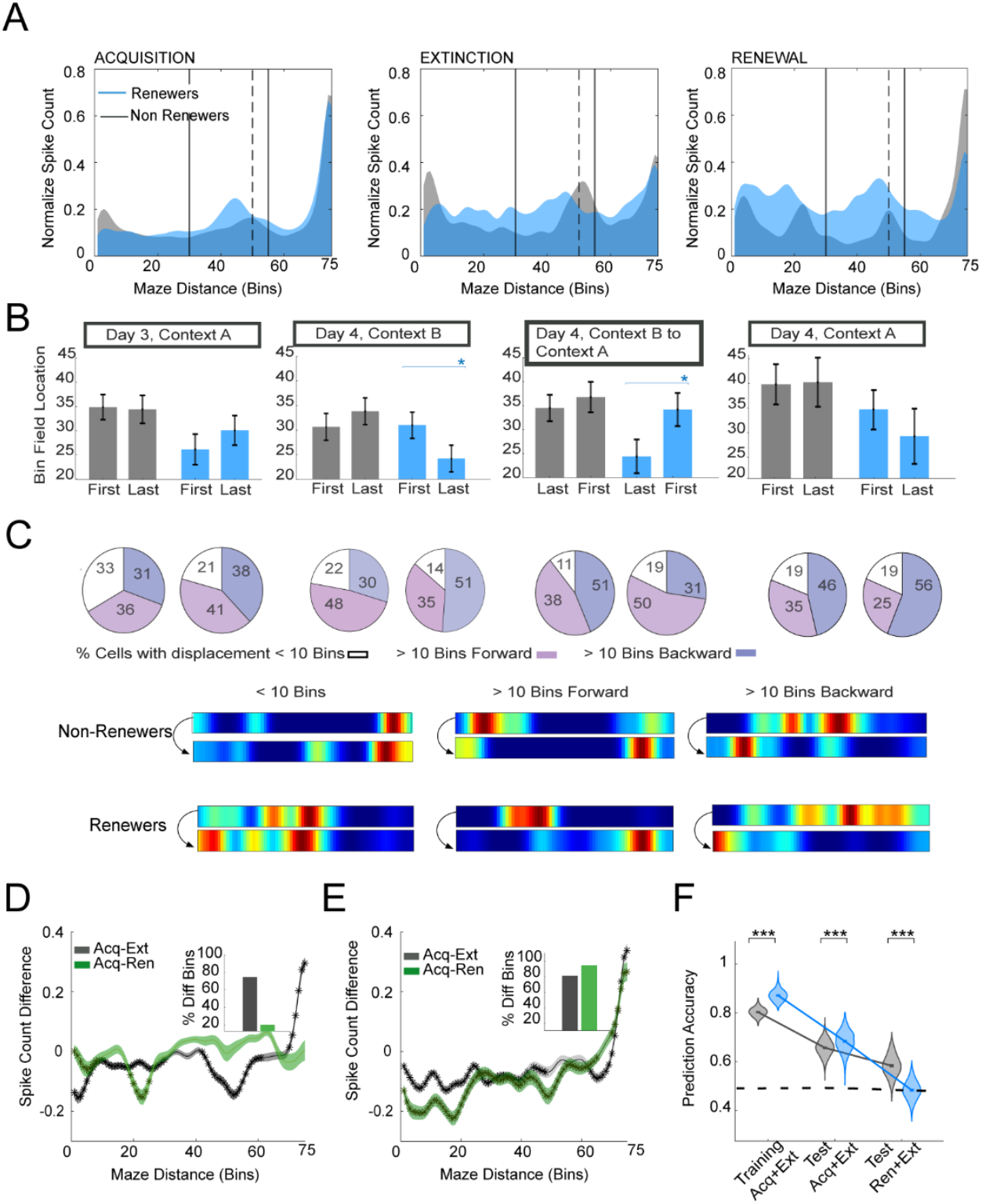
Renewers exhibit more context-dependent place cell remapping. **(A)** Normalized spike count distributions across the maze for renewers (blue) and non-renewers (grey) across experimental phases. **(B)** Mean place field bin locations across trial transitions, showing context-dependent shifts in renewers but stable field positions in non-renewers. Error bars represent 95% CI. **(C)** Top: Place field displacement categorization (<10 bins, >10 bins forward or backward). Spatial arrangement of pie charts according to that of bars in B. Bottom: examples of cell in the three categories for the two groups. Renewers exhibited more pronounced forward or backward shifts aligned with context changes. (**D-E**) Spike count differences by bin for non-renewers (D) and renewers (E), comparing Acquisition (Acq) vs. Extinction Learning (Ext, gray) and Acquisition vs. Renewal testing (Ren, green). Bar plots show the proportion of bins differing by Kolmogorov–Smirnov tests (*p<0.05). (**F**) Violin plot representing predicted accuracies for the ML models (1000 randomised iterations), where Training Acq+Ext represents the predicted accuracies for the training data set; Test Acq+Ext represents the accuracies for the model containing the test data of acquisition and Extinction Learning; and Test Ren+Ext for the model containing the test data for renewal and Extinction Learning. Models for renewers failed to classify renewal trials, consistent with remapping, whereas non-renewers showed above-chance performance.

We next explored global remapping through assessment of PC displacements (Fig. 3B; *SI Appendix* Table S6). In renewers, peak field locations systematically shifted forward in context A (acquisition and renewal) and backward in context B (EL), consistent with context-sensitive remapping. In contrast, non-renewers showed stable field locations across all phases (Fig. 3B). Despite similar proportions of small vs. large shifts (<10 or >10 bins; Fig. 3C), only renewers displayed context-dependent directional changes, indicating flexible spatial coding. Total counts of fields in the maze supported these results (*SI Appendix* Fig. S3B and Table S7). Spatial correlations subtracted across acquisition, EL and renewal phases further supported these findings (*SI Appendix* Fig. S3C and Table S7): correlations in renewers from EL to renewal dropped significantly in comparison with those correlations within contexts, reflecting distinct context-related information processing. This effect was not observed in non-renewers.

To further assess the spatial reorganization of PC responses in renewers and non-renewers, we compared their spike count distributions at different locations in the maze between acquisition (day 3) and either EL, or renewal (Fig. 3D-E). During EL, about 80% of the bins differed from acquisition in both groups; however, the pattern diverged: non-renewers (Fig.3D) clustered spikes in the DZ, while renewers (Fig. 3E) displayed a wider spread of spikes throughout the T-maze. During renewal, spike distributions in non-renewers were similar to those observed in acquisition, with only 20% of the bins differing. In contrast, renewers showed significant changes in 92% of the bins, which is consistent with the remapping and re-encoding of context A after EL in context B. Logistic regression classifiers (trained on spike distributions) confirmed context-specific remapping (Fig. 3F; SI Appendix Table S7): Renewers distinguished contexts A and B during acquisition and EL (AUC = 0.871), but failed to classify renewal trials correctly as reflected by remapping (below-chance accuracy, 0.484), unlike non-renewers (0.582), that showed stable spatial maps.

In sum these results indicate that PCs in renewers undergo dynamic remapping in response to changes in context and reward contingencies, and that these processes support behavioral context-dependent renewal. In contrast, the stable and context-invariant activity of PCs in non-renewers across acquisition, EL and renewal testing suggest reduced hippocampal flexibility, possibly due to the absence of remapping. These findings align with computational models proposing global remapping as a neural mechanism for behavioural renewal (16).

## Discussion

Whereas an absence of renewal (behavioral re-activation) may be a desirable outcome of extinction learning (EL) of aversive experience, for more benign forms, learning related to associative daily experience, renewal may be of value. For instance, this could reflect the resumption of an individual to adopt a spatial route (that had been abandoned in winter) to an outdoor market to buy fruit that is once more in season. For rodents, this might encompass the resumption of the search for food remnants under a park bench on a sunny day (context A), having learned that this is unlikely to occur when it rains (context B). Even in very standardized spatial appetitive learning paradigms, not all rodents exhibit effective renewal, despite showing effective context-dependent task acquisition and EL (17, 18). By means of place cell recordings conducted during acquisition, EL and renewal testing in a spatial appetitive T-maze task, we show here that the different task acquisition and EL strategies adopted by rats determine the outcome of renewal. Our previously established computational model (16) validated this interpretation and demonstrated, moreover, that differentiation of contextual representations during EL is a prerequisite for renewal to occur. In line with the model’s prediction, in our experiment the extent of global remapping of spatial representations predicts whether context-dependent renewal will succeed or fail. These findings provide important insights into context-dependent learned behavior.

Our behavioral paradigm was designed so that acquisition learning was cumulative, effortful and dependent on spatial cues (10). Despite earlier assumptions that spatial (i.e. declarative) and non-declarative learning depend on distinct brain structures and mechanisms that largely do not overlap (19), more recently, evidence has accumulated that an interaction of these memory systems supports hippocampus-dependent learning (20). Moreover, neuromodulatory systems such as the dopaminergic ventral tegmental area (VTA) potently influence both hippocampal information processing and storage (9, 21), as well as non-declarative forms of learning (22, 23).

Although both acquisition learning and EL were ostensibly similar in animals that later emerged as renewers or non-renewers, their behavior during acquisition training revealed subtle differences. For example, renewers left the start-box faster than non-renewers and spent less time in the T-maze intersection and junction segments before deciding on which arm to enter. Single-unit recordings revealed higher power of theta frequency oscillations during correct arm choices in renewers, as well as tighter organization of place cell firing within theta cycles, compared to non-renewers, consistent with a higher efficacy of spatial learning (24, 25). In general, place cells of renewers exhibited higher peak firing rates than renewers during acquisition, EL and renewal. These differences may have derived to some extent from different speeds of movement of renewers and non-renewers in the different segments of maze (26), especially given that place cell burst firing properties were identical in the two groups indicating that recordings were made from the same sublayers in CA1 (27). However, renewers showed increased peak firing during EL compared to acquisition, engaged in more effective spatial encoding of the T-maze contexts (11). Furthermore, examination of place fields revealed that during acquisition, renewers exhibited a predominance of place fields in the decision zone, whereas non-renewers showed more fields in the stating box of the T-Maze, possibly reflecting initially different learning proficiencies (28). Moreover, renewers exhibited rate remapping during EL, consistent with an updating of non-spatial information (29) and task adjustment (30). Renewers also exhibited more global remapping, consistent with their perception of the change in context. These findings may be consistent with recent findings that different neurons within CA1 may be responsible for the encoding and updating of spatial and non-spatial elements of a spatial representation (31). By contrast, non-renewers exhibited stable place cell behavior when activity was compared between acquisition and EL, suggesting that the change of context or indeed informational content, was not perceived by the animals as relevant to the task.

If context was a driving factor in the differences of place cell behavior in renewers compared to non-renewers, one would (in addition to abovementioned remapping during EL) expect to see further remapping in renewers during renewal testing. In contrast to non-renewers, place cells in renewers exhibited increased firing compared to their own activity during acquisition, as well as a degradation of place field representations (spatial information scores), properties which may reflect greater spatial flexibility (32, 33). This suggests that in renewers, information updating occurred as the animal realized that context no longer contained a reward (18). Non-renewers exhibited spatial properties that were similar to those detected during acquisition. This suggests that non-renewers did not attend to the context and that it was the attention to the reward location that motivated both EL and failure of renewal in this group (32), akin to behavioral profiles when EL is tested in an AAA paradigm (4).

If this is the case, the place cell activity during EL may offer neural signatures of learning that impact on renewal efficacy. To test this we used logistic regression analysis and confirmed that place cell behavior during EL could predict whether the animals would re-instate their previously learned behavior, when returned to context A. Place cell firing within a spatial environment can anchor to high-valence information (34, 35). We detected a greater degree of spiking activity in the decision zone of renewers during acquisition trials, whereas non-renewers exhibited more firing activity in the starting box. In renewers the concentration point of spikes around the decision zone matched their maximum deceleration moment, indicating that renewers engaged in cognitive, spatially driven, decision-making (36). Strikingly, during EL, non-renewers showed more spiking activity in the decision zone despite an absence of place field remapping, whereas renewers showed shifts of firing across the T-maze (as reflected by the global remapping described above). This might reflect differences in the state-dependent responses of the hippocampus to the absence of the reward (9, 21, 37). This, in turn, may correspond to known intra-individual differences in rats with regard to their response to cue motivation (i.e. interpretation of cues, as incentive versus defeat-related) (38).

In conclusion, this study provides neurophysiological and behavioral evidence that the precise nature of the information acquisition/updating strategy, during acquisition learning and EL of spatial appetitive experience, determines renewal efficacy. Our data indicate that that global remapping of place fields during context-dependent EL serves as a key neural mechanism underlying effective renewal. This process allows for the segregation of distinct memories and the prevention the generalization of EL with initial learning experiences. Moreover, our data indicate that place cells may serve to integrate both spatial and non-spatial information during associative learning and the updating of learned spatial experience.

## Supporting information

Supplementary methods, figures and tablet

## Materials and methods

### For complete Materials and Methods, see SI Appendix, Materials and Methods

#### Animals

Thirteen male Long-Evans rats (8–12 weeks old) were housed under standard conditions with ad libitum access to water and food. After handling habituation, rats were implanted with movable tetrodes (Axona Ltd, St. Albans, UK) targeting dorsal CA1 (AP −3.8, ML ±2.5, DV −2.5 mm), as previously described (Dolón et al., 2024). Experiments were conducted in accordance with EU Directive 2010/63 and approved by NRW authorities. Rats were food-restricted to 85% baseline before training.

#### ABA paradigm

Animals participated in a 4-day-long spatial appetitive ABA learning paradigm in a T-maze, consisting of three phases (6): acquisition, extinction learning (EL) and renewal. During acquisition (days 1–3), rats learned to collect a reward at the end of the right arm over 20 trials/day, with reward probability gradually reduced from 100% on day 1 to 30% on day 3. On day 4, EL was conducted in context B (15 unrewarded trials), immediately followed by renewal testing in context A (5 unrewarded trials). Context A and B diverged in distal cues, floor patterns and odors. Correct responses were defined as entries into the rewarded arm. Acquisition was defined as >80% correct responses on last 5 trials of day 3. Three animals did not meet acquisition criteria and were excluded. EL was defined as a decrease of <70% correct responses from the last 5 trials of acquisition to the last 5 trials of EL. Renewal was defined as >80% correct responses in 5 renewal trials.

#### Data Collection & analyses

Neuronal and behavioral data were acquired using the dacqUSB system (Axona Ltd., St. Albans, UK). Unless noted, analyses were performed in MATLAB using custom scripts (MATLAB 2021a and later, Mathworks, USA).

**Animal position** was tracked at 50 Hz and linearised into 150 bins (1cm/bin). Speed was calculated as Euclidean distance between consecutive points divided by the time interval (20ms); angular velocity was calculated from the cross product of position and velocity vectors to quantify turning speed (radians/s).

**Single-unit activity** was sampled at 48 kHz, amplified 10000-30000 times, and band-pass filtered (600-7000 Hz). Spike sorting was performed offline using TINT software and in-program Klustakwik algorithm (Axona Ltd, St. Albans, UK). Putative interneurons (waveform’s width from peak to valley <200µs and average firing rates >10Hz) were excluded (39). Spike counts and occupancy maps of 75 bins were generated by associating spike times with the temporally closest linearized position (2cm/bin), and smoothed firing rate maps were constructed by dividing the spike count map by the occupancy map with a smoothing factor of 2 bins. Place fields were then defined based on the peak firing rate (maximum value in the rate map) and their field width (sum of the number of contiguous active bins where a neuron’s firing rate exceeds 20% of its peak rate). Place field features were extracted to quantify spatial coding (40). Average firing rates were calculated by dividing the number of spikes that occurred over the entire session by the total duration of the session; infield firing rates were calculated as the mean firing rate for all bins within the field; outfield firing rate as the mean firing rate for all bins outside of the field; percentage of active bins as the sum of all bins in which a neuron was active exceeding the 50% of the peak firing rate; spatial information content, comprising the amount of information (in bits) conveyed about the spatial location of the animal by a single spike from a cell, was calculated using the method described by Skaggs and colleagues (41); and information per second (bits/s), defined as the rate at which a neuron transmits information, was calculated as described previously (42). Finally, spatial coherence, which is a measure of how spatially contiguous the neuron’s activity is, was calculated as described by Valero and colleagues (40) by subtracting the mean Pearson correlation between the firing rate in each spatial bin and the corresponding rates averaged over the ±4 nearest-neighbor bins (calculated using a 3×3 filter kernel). Place cells were defined as cells meeting the following criteria: spatial information content score >= 0.50, (43, 44), field size >6 bins (40), spike waveform width from peak to valley >200µs and average firing rates between 0.1 and 10Hz (39).

**Local field potentials** (LFPs) were notch-filtered (50 Hz), amplified (1000–5000×), sampled at 4800 Hz. Theta phase was derived from the Hilbert transform (5–11 Hz), and theta epochs identified via the ratio of the power in theta band (5–11 Hz) to the power of nearby bands (1–4 Hz, 12–14 Hz) (40, 45). Phase locking was assessed by mean resultant length; only cells with significant Rayleigh tests (p<0.05) were included. Adapted code from public repository HippoCookBook (GitHub - valegarman/HippoCookBook: MATLAB-based repository for electrophysiological experiments and data analysis).

#### Logistic regression models

To assess predictors of renewal, we performed a logistic GLM using renewal expression (yes/no) as the outcome and place field features across days (peak and average firing rates, infield and out/in firing rates, spatial information, coherence, bits/s, field size) as predictors. The model was fitted in MATLAB (fitglm), using a binomial distribution and logit link. Predicted probabilities were used to compute ROC curves and AUC to quantify discrimination, and model coefficients identified key spatial features associated with renewal. For context decoding, two logistic regression models were implemented (adapted from Dr. Milan Parmar’s “Machine Learning Algorithms from Scratch”). These models classified contexts (A vs. B) using normalized binned spike counts from PCs. Both employed Iterative Reweighted Least Squares. The first model was trained on day 1 acquisition (context A) and EL (context B) data (70%) and tested on the remaining 30% trials; the second was tested on renewal trials versus extinction. Each model was run for 1000 independent iterations per group to derive classification metrics (accuracy, precision, recall, AUC).

#### Statistical analysis

Significance was set at p < 0.05. Normality was assessed using the Kolmogorov-Smirnov test. One-way or two-way rmANOVA was used to evaluate differences between sessions and groups, followed by Bonferroni-corrected *post-hoc* tests. Non-normal data were compared using Wilcoxon rank-sum tests; two-sample Kolmogorov-Smirnov tests assessed distribution differences. Behavioral response proportions were analyzed with chi-square tests. Circular data (theta phase modulation) were assessed using the Watson-Williams multi-sample test (CircStat toolbox). Effect sizes were calculated as Cohen’s d.

## Acknowledgments

This study was supported by a grant from the German Research Foundation (Deutsche Forschungsgemeinschaft) through SFB 1280 (project number: 316803389) to DMV (subproject A04) and SC (subprojects A14, F01). We thank Juliane Böge and Ute Neubacher for technical assistance, Dr. Marta Méndez-Couz for support with behavioral experiments, and Nadine Kollosch for animal care.

## References

1. Z. Wen, Z. S. Chen, M. R. Milad, Fear extinction learning modulates large-scale brain connectivity. NeuroImage 238, 118261 (2021).

2. L. Pessoa, How many brain regions are needed to elucidate the neural bases of fear and anxiety? Neurosci. Biobehav. Rev. 146, 105039 (2023).

3. M. E. Bouton, S. Maren, G. P. McNally, Behavioral and neurobiological mechanisms of pavlovian and instrumental extinction learning. Physiol. Rev. 101, 611–681 (2021).

4. M. A. E. André, O. T. Wolf, D. Manahan-Vaughan, Beta-adrenergic receptors support attention to extinction learning that occurs in the absence, but not the presence, of a context change. Front. Behav. Neurosci. 9 (2015).

5. S. Trent, et al., Fear conditioning: Insights into learning, memory and extinction and its relevance to clinical disorders. Prog. Neuropsychopharmacol. Biol. Psychiatry 138, 111310 (2025).

6. M. Méndez-Couz, J. M. Becker, D. Manahan-Vaughan, Functional Compartmentalization of the Contribution of Hippocampal Subfields to Context-Dependent Extinction Learning. Front. Behav. Neurosci. 13 (2019).

7. H. Eichenbaum, The role of the hippocampus in navigation is memory. J. Neurophysiol. 117, 1785–1796 (2017).

8. J. E. Lisman, A. A. Grace, The hippocampal-VTA loop: controlling the entry of information into long-term memory. Neuron 46, 703–713 (2005).

9. H. Hagena, D. Manahan-Vaughan, Oppositional and competitive instigation of hippocampal synaptic plasticity by the VTA and locus coeruleus. Proc. Natl. Acad. Sci. 122, e2402356122 (2025).

10. V. Wiescholleck, M. A. Emma André, D. Manahan-Vaughan, Early age-dependent impairments of context-dependent extinction learning, object recognition, and object-place learning occur in rats. Hippocampus 24, 270–279 (2014).

11. K. J. Jeffery, Integration of the sensory inputs to place cells: What, where, why, and how? Hippocampus 17, 775–785 (2007).

12. Y. Aoki, H. Igata, Y. Ikegaya, T. Sasaki, The Integration of Goal-Directed Signals onto Spatial Maps of Hippocampal Place Cells. Cell Rep. 27, 1516-1527.e5 (2019).

13. Z. Navratilova, L. T. Hoang, C. D. Schwindel, M. Tatsuno, B. L. McNaughton, Experience-dependent firing rate remapping generates directional selectivity in hippocampal place cells. Front. Neural Circuits 6, 6 (2012).

14. S. Krishnan, C. Heer, C. Cherian, M. E. J. Sheffield, Reward expectation extinction restructures and degrades CA1 spatial maps through loss of a dopaminergic reward proximity signal. Nat. Commun. 13, 6662 (2022).

15. C. B. Alme, et al., Place cells in the hippocampus: Eleven maps for eleven rooms. Proc. Natl. Acad. Sci. 111, 18428–18435 (2014).

16. D. Kappel, S. Cheng, Global remapping emerges as the mechanism for renewal of context-dependent behavior in a reinforcement learning model. [Preprint] (2023). Available at: https://www.biorxiv.org/content/10.1101/2023.10.27.564433v1 [Accessed 29 July 2024].

17. P. Gasalla, D. Manahan-Vaughan, D. M. Dwyer, J. Hall, M. Méndez-Couz, Characterisation of the neural basis underlying appetitive extinction & renewal in Cacna1c rats. Neuropharmacology 227, 109444 (2023).

18. J. Haubrich, L. D. Vera, D. Manahan-Vaughan, Cortico-subcortical networks that determine behavioral memory renewal are redefined by noradrenergic neuromodulation. Sci. Rep. 15, 9692 (2025).

19. B. Milner, L. R. Squire, E. R. Kandel, Cognitive Neuroscience and the Study of Memory. Neuron 20, 445–468 (1998).

20. J. M. Froula, S. D. Hastings, E. Krook-Magnuson, The little brain and the seahorse: Cerebellar-hippocampal interactions. Front. Syst. Neurosci. 17 (2023).

21. O. Mamad, et al., Place field assembly distribution encodes preferred locations. PLOS Biol. 15, e2002365 (2017).

22. I. Carta, C. H. Chen, A. L. Schott, S. Dorizan, K. Khodakhah, Cerebellar modulation of the reward circuitry and social behavior. Science 363, eaav0581 (2019).

23. M.-Y. Jing, et al., Re-examining the role of ventral tegmental area dopaminergic neurons in motor activity and reinforcement by chemogenetic and optogenetic manipulation in mice. Metab. Brain Dis. 34, 1421–1430 (2019).

24. M. C. Zielinski, J. D. Shin, S. P. Jadhav, “Coherent coding of spatial position mediated by theta oscillations in hippocampus and prefrontal cortex” (Neuroscience, 2019).

25. B. E. Pfeiffer, Spatial Learning Drives Rapid Goal Representation in Hippocampal Ripples without Place Field Accumulation or Goal-Oriented Theta Sequences. J. Neurosci. 42, 3975–3988 (2022).

26. A. P. Maurer, S. R. Vanrhoads, G. R. Sutherland, P. Lipa, B. L. McNaughton, Self-motion and the origin of differential spatial scaling along the septo-temporal axis of the hippocampus. Hippocampus 15, 841–852 (2005).

27. K. Mizuseki, K. Diba, E. Pastalkova, G. Buzsáki, Hippocampal CA1 pyramidal cells form functionally distinct sublayers. Nat. Neurosci. 14, 1174–1181 (2011).

28. A. Rayan, et al., Learning shifts the preferred theta phase of gamma oscillations in CA1. Hippocampus 32, 695–704 (2022).

29. H. Sanders, et al., Temporal coding and rate remapping: Representation of nonspatial information in the hippocampus. Hippocampus 29, 111–127 (2019).

30. K. Allen, J. N. P. Rawlins, D. M. Bannerman, J. Csicsvari, Hippocampal Place Cells Can Encode Multiple Trial-Dependent Features through Rate Remapping. J. Neurosci. 32, 14752–14766 (2012).

31. T-H. Hoang, D. Manahan-Vaughan, Multiplexed detection of nuclear immediate early gene expression reveals hippocampal neuronal subpopulations that engage in the acquisition and updating of spatial experience. BIORXIV/2025/663345 (2025).

32. S. Krishnan, M. E. J. Sheffield, Reward Expectation Reduces Representational Drift in the Hippocampus. [Preprint] (2023). Available at: https://www.biorxiv.org/content/10.1101/2023.12.21.572809v1 [Accessed 27 July 2024].

33. Y. Chiu, C. Dong, S. Krishnan, M. E. J. Sheffield, The Precision of Place Fields Governs Their Fate across Epochs of Experience. eNeuro 10 (2023).

34. R. E. Ambrose, B. E. Pfeiffer, D. J. Foster, Reverse Replay of Hippocampal Place Cells Is Uniquely Modulated by Changing Reward. Neuron 91, 1124–1136 (2016).

35. J. L. Gauthier, D. W. Tank, A Dedicated Population for Reward Coding in the Hippocampus. Neuron 99, 179-193.e7 (2018).

36. U. Mugan, S. L. Hoffman, A. D. Redish, Environmental complexity modulates information processing and the balance between decision-making systems. Neuron 112, 4096-4114.e10 (2024).

37. A. M. Kaufman, T. Geiller, A. Losonczy, A Role for the Locus Coeruleus in Hippocampal CA1 Place Cell Reorganization during Spatial Reward Learning. Neuron 105, 1018-1026.e4 (2020).

38. T. E. Robinson, L. M. Yager, E. S. Cogan, B. T. Saunders, On the motivational properties of reward cues: Individual differences. Neuropharmacology 76 Pt B, 450–459 (2014).

39. R. E. Harvey, et al., Linear Self-Motion Cues Support the Spatial Distribution and Stability of Hippocampal Place Cells. Curr. Biol. 28, 1803-1810.e5 (2018).

40. M. Valero, A. Navas-Olive, L. M. De La Prida, G. Buzsáki, Inhibitory conductance controls place field dynamics in the hippocampus. Cell Rep. 40, 111232 (2022).

41. W. E. Skaggs, B. L. McNaughton, K. M. Gothard, E. J. Markus, An Information-Theoretic Approach to Deciphering the Hippocampal Code in In, (Morgan Kaufmann, 1993), pp. 1030–1037.

42. B. C. Souza, R. Pavão, H. Belchior, A. B. L. Tort, On Information Metrics for Spatial Coding. Neuroscience 375, 62–73 (2018).

43. É. Duvelle, et al., Insensitivity of Place Cells to the Value of Spatial Goals in a Two-Choice Flexible Navigation Task. J. Neurosci. 39, 2522–2541 (2019).

44. R. M. Grieves, E. R. Wood, P. A. Dudchenko, Place cells on a maze encode routes rather than destinations. eLife 5, e15986 (2016).

45. A. Fernández-Ruiz, et al., Entorhinal-CA3 Dual-Input Control of Spike Timing in the Hippocampus by Theta-Gamma Coupling. Neuron 93, 1213-1226.e5 (2017).

