## Supplementary methods, figures and tablet for "Place cell activity and behaviour during task acquisition and extinction learning predict renewal outcome"

**This PDF file includes:**

Supplementary material and methods
Figures S1 to S3
Tables S1 to S7
SI References

**Other supporting materials for this manuscript include the following:**

### Supporting Information Text

#### DETAILED MATERIAL AND METHODS

**Animals** were housed in polypropylene cages on a 12-h light/dark cycle, with ad-libitum access to water and food. Cages were stored in a vivarium (Scantainer, Scanbur Technology A/S, Denmark) with humidity and temperature control (45-50% humidity, 21-23°C). Before any experimental procedure, rats were housed with their cage mates and handled daily for habituation. Upon reaching 8 to 12 weeks of age, they underwent surgery, housed individually, weighed and handled daily to ensure they were in good health. Before the behavioral protocol started, rats were food deprived to 85% of their initial body weight.

**Surgery and Electrode Preparation**:

Male Long Evans Rats (8 to 12 weeks old) were chronically implanted with a custom-built 32-channel drive in the CA1 area of the dorsal hippocampus as previously described (1). The microdrives used in this study were lightweight microdrives (Axona Ltd, St. Albans, UK) and held 8 tetrodes. Tetrodes were created using 25 µm diameter formvar-coated platinum-iridium wires (A-M systems, Sequim, WA, USA) and a tetrode assembly station (Neuralynx, Bozeman, MT, USA). The tetrode tips were electroplated to reduce impedance to around 250 kΩ (Ferguson et al., 2009). The electroplating was performed in gold solution (Gold Plating Solution, Axona Ltd, St Albans, UK) using an impedance meter (nano Z, Neuralynx, Bozeman, MT, USA) and by passing negative current throughout each wire.

Animals were deeply anaesthetized with sodium pentobarbital (52mg/kg intraperitoneally) and placed in a stereotaxic frame (430005-series, TSE Systems, Bad Homburg, Germany) for tetrode implantation as previously described (1). The analgesic, meloxicam (0.2mg/kg, i.p.), was given before surgery and both 24 h and 48 h afterwards. The tetrode was implanted 3.8mm anteroposterior and 3.0mm mediolateral from bregma (2). A craniotomy (1.2 mm diameter) was made, through which the microdrive was carefully inserted. It was slowly lowered until the tetrodes were placed in the cortex with their tips above the CA1 region (2.5 mm dorsoventral from the skull). The microdrive was anchored to the skull with fixation screws and dental acrylic (Paladur, Heraeus Kulzer GmbH). Once the microdrive was fixed, the guiding gauge for the tetrode was lowered. The space between the skull and the gauge was sealed with vaseline (ISANA, Dirk Rossmann GmbH) and the rest of the surgical field was covered with dental acrylic. After surgery, rats recovered for at least 10 days before any screening started and their health was monitored daily.

**Behavioral apparatus and protocols:** Handling habituation was performed daily for 10 to 20 minutes over a 3 week period before surgery. A week before surgery, rats were habituated to a square arena (80 cm x 80 cm grey polyvinylchloride, PVC, with 70 cm high walls) with their cage mates for two days (30 min/day), which was later used for recordings to target the CA1 area during free foraging (30min/day). Chocolate sprinkles (ca. 4 x 1mm, Dr. Oetker, Germany) were used as a movement incentive during all behavioral procedures. When stable single units were detected, animals participated in the ABA paradigm in a T-maze based on previous protocols (3) (Figure 1A).

The maze was made of grey PVC and consisted of a starting box (**S**, 25 × 20 cm), a corridor (100 × 20 cm), a sliding door that separates the starting box from the corridor, and two perpendicular arms (12 × 40 cm each) (**SI** Appendix Fig. S1) with a small circular indentation in the floor at the end of these arms that served to conceal detection of the reward from the T-maze junction.. The T-maze (and abovementioned square arena) were enclosed in a square-shaped recording tent (3x3m) made of black fabric for acoustic insulation (Tolko Stoffe GmbH, Germany) and a pure-silver-metallized polyamide fabric for electromagnetic insulation (Shieldex® Bremen PW). The tent was equipped with controlled LED illumination. The behavioral protocol consisted of four phases: habituation (4 days), acquisition (3 days), EL (day 4, sessions 1–3), and renewal (day 4, session 4). Each experimental day consisted of 4 sessions separated by a resting period of 10 minutes, during which rats were removed from the maze and allowed to rest in a resting box. Each session consisted of 5 trials separated by 15s in **S**. This time was also used for cleaning the maze and placing the reward. No floor patterns, visual cues, or scents were used during habituation.

Two contexts were created: Context A (acquisition and renewal) featured a granite floor pattern, a black circle visual cue, white LED light, and vanilla scent; while Context B (EL) a wood floor pattern, a white circle visual cue, warm yellow LED light, and almond scent. No context change was created for **S**, i.e. this area was always constant in appearance.

Habituation Day 1 consisted of 5 min of free exploration with randomly scattered chocolate sprinkles throughout the T-maze. On Day 2, rats waited in **S** for 15s and freely explored the T-maze during 2 min with randomly scattered sprinkles in the arms only. On Days 3-4, animals performed 2 sessions of 5 trials each with sprinkles only at the reward indentations in the T-maze arms. Here, animals were allowed to explore for 2 min or until they found the sprinkles and then guided back to S. During acquisition trials, one T-maze arm was rewarded, and reward probability was gradually reduced from 100% on Day 1 to 30% in the final session of Day 3. The reward (if present) was always located at the right arm, and its location was consistent for all animals.

**Data Collection:** Neuronal and behavioural data were acquired using the dacqUSB system (Axona Ltd., St. Albans, UK). Signals were passed through AC-coupled, unity-gain headstage amplifiers connected near the rat's head and then fed into a preamplifier (Axona Ltd, St. Albans, UK). One tetrode channel was used for LFP recordings, and the remaining 31 channels recorded single-unit activity. Single-unit activity was sampled at 48 kHz, amplified 10000-30000 times, and band-pass filtered between 600-7000 Hz. Spikes were stored in 1ms windows (200 µs pre-trigger, 800 µs post-trigger) per tetrode group, and each tetrode was referenced by one electrode from another tetrode. LFPs were notch-filtered (50 Hz notch filter), amplified (1000-5000 times) and sampled at 4800 Hz. Animal position was tracked at 50Hz with a monochrome video camera (CAM-M1, Axona Ltd, St. Albans, UK) and the data was converted into x-y coordinates using Axona's software. Following recovery, rats were screened daily, five days a week, in the abovementioned square arena (30 min). These recordings aimed to target the dorsal CA1 pyramidal layer and find place cells. Single-unit activity was examined offline using TINT software (Axona Ltd, St. Alban, UK), and local field potentials (LFPs) were analysed using MATLAB (Mathworks, USA). Tetrodes were advanced 25-50µm daily until stable single-unit activity was detected, with a maximum movement of 100µm per day. The experiments started once single-unit activity was identified. Tetrodes were slightly retracted after each experimental day to preserve the neurons overnight, and were then relocated on the next experimental day.

**Spike Sorting and Cell Matching:** Spike sorting was performed offline using TINT software (Axona Ltd, St. Albans, UK) and the Klustakwik algorithm. For each tetrode, spike waveforms dimensionality was reduced to the first principal component, peak-trough distance, voltage at time, energy, peak height, and peak time. Klustakwik was then applied to isolate waveforms into clusters representing putative neurons, which were further checked, combined, or refined manually. Only clusters with more than 50 spikes and without 2ms refractory period violations were included (1). Clusters belonging to the same putative neuron recorded in different experimental sessions were identified using Multisession Cut File Splitter software (Axona Ltd, St. Albans, UK).

**Classification of behavioral responses:** Behavioral responses were categorized as Correct (animal departed from **S** within 50s and visited the rewarded arm), Incorrect (animal departed from **S** within 50s and visited the left arm), Back (animal departed from **S** within 50s and turned back before entering any arm or remained static for a period longer than 50s), or Indecision (animal did not leave **S** within 50s). When a rat entered the left arm, it was blocked with a barrier for 15s before being allowed to return to **S**. Animals scoring less than 80% Correct responses in the final session of Day 3 Acquisition were excluded from the study. Three animals were excluded. Given that each session consisted of five trials, the probability of an animal performing three correct trials by chance is less than 18.75% (calculated as the binomial probability of three or more successes out of five trials with $p = 0.5$). Extinction learning (EL) was considered successful when animals exhibited a 70% reduction in correct arm choices from the final acquisition session to the final EL trial. Renewal was defined as an increase of 80% or more in correct responses from the final EL session to the end of the renewal session.

**Standardisation and linearisation of the animals' position:** X and y position coordinates were normalised by rescaling their values. Y coordinates range between -0.2 and 1.2, where the negative values represent S, values between 0 to 0.7 in the corridor (C), and from 0.71 to 1.2, the junction (**J**) and intersection (**I**) zones (Figure 1E). X coordinates range from -1 to 1, where negative values represent the left arm of the maze and positive values represent the right arm. Then, the two-dimensional position of the rats was linearized according to the method described by van der Meer and Redish (4).

**Cell activity maps and basic PCs parameters:** One-dimensional rate maps were generated with adapted code from the open-source repository HippoCookBook (GitHub -

valegarman/HippoCookBook: MATLAB-based repository for electrophysiological experiments and data analysis) created by Manuel Valero. First, the linearized position was binned into 2 cm wide bins (75 bins for the totality of the maze), and the time of occurrence of each spike was associated with the temporally closest recorded position for each bin. Then, spike counts and occupancy maps of 75 bins were generated. After, smoothed firing rate maps were constructed by dividing the spike count map by the occupancy map with a smoothing factor of 2 bins. The same smoothing parameter was applied to all rate maps. Place fields were then defined based on the peak firing rate (maximum value in the rate map) and their field width (sum of the number of contiguous active bins where a neuron's firing rate exceeds 20% of its peak rate). Only the main field containing the peak firing rate was included in the analyses. Other spatial modulation measurements were calculated as previously described (5, 6).

**Speed assessment and speed maps:** The linearized positions were used to calculate running speed (as Euclidean distance between consecutive points sampled at 50 Hz) and angular velocity (as the cross product of position and velocity vectors, in radians/s). Periods <2 cm/s were excluded. Speed and angular velocity were then binned into 1 cm-wide spatial bins (150 bins) to generate maps across the maze.

**Spike Shuffling and Cell criteria inclusion:** To ensure that only place-selective cells were included in the analyses, a first exclusion criterion was created based on comparing the spatial information scores of cells with an artificial shuffled distribution. For each cell and trial, spike times were temporally shifted by a random interval ranging from 10 seconds to the total trial length minus 10 seconds. Subsequently, these shifted spike times were utilized to construct firing rate maps, from which the spatial information score was computed. This process was iterated 1000 times for each cell and trial, resulting in a distribution of spatial information scores. The obtained distribution of spatial information scores was then compared to the original information score derived from the unshifted spike times. Only cells, the spatial information score of which exceeded the 95th percentile of this shuffled distribution, were included in the analyses. Later, for each cell and trial, putative neurons were classified as place cells if they met the following criteria: spatial information content score >= 0.50, (7, 8), field size bigger than 6 bins (5), waveform's width from peak to valley >200μs and average firing rates <10Hz but higher than 0.1 Hz (9). Only PCs that fulfilled these criteria were included in the analyses. Fast spiking putative neurons were classified based on the waveform's width from peak to valley <200μs and high average firing rates (>10Hz) (9) and excluded from all analyses.

**Power Spectrum analyses:** LFP data was high-pass filtered at 4 Hz and notch filtering at 50 Hz, and down-sampled to 1200 Hz using a Finite Impulse Response (FIR). Power spectral density was estimated via Welch's method using the *pwelch* function in MATLAB. All trials for each animal and day were concatenated to construct "hyper-trials" with zero padding of 0.5 seconds on the edges of each trial. Then, a Hamming window of 2 seconds and a frequency resolution of 100 times the sampling rate were used as parameters for the welch function. The overlap between consecutive segments was 25% of the window size.

**Spike lock to theta:** Theta modulation indices for each neuron were estimated with adapted code from the public repository HippoCookBook in GitHub from Dr. Manuel Valero (mentioned above) and as described previously by Fernández-Ruiz et al. (10). First, the theta-band phase of the LFP was estimated as the Hilbert transform of the narrowband filtered LFP (5–11 Hz). Then, theta epochs were detected automatically using the ratio of the power in theta band (5–11 Hz) to the power of nearby bands (1–4 Hz, 12–14 Hz) (5, 10). Theta peaks correspond 0 rad (0 deg) and 2pi rad (360 deg) and troughs at pi rad (180 deg). Last, the theta indices for each neuron were extracted by calculating the mean resultant length of the phases, and the significance of the modulation was estimated with the Rayleigh test for non-uniformity. Only cells with significant Rayleigh scores ($p<0.05$) were included in the analysis. The mean angle and mean resultant length of the theta phases for each neuron were taken as the preferred phase and modulation strength of that neuron.

**GLM for renewal expression prediction:** a logistic regression analysis (logistic GLM) was conducted to assess how predictive PCs spatial properties were for the expression of renewal. The outcome variable was the presence of renewal (0 = no expression of renewal; 1 = expression of renewal). Predictor variables included experimental days (day 1, day 2, day 3, and day 4, but only trials from EL) and basic place field features. The GLM was fitted using the MATLAB function *fitglm*. The model equation was: *glmModel* =

*fitglm(T, 'Group ~ AV*Day + Peak*Day + Bits*Day + SI*Day + SC*Day + size*Day + Outratio*Day + Infield*Day', 'Distribution', 'binomial','Link','logit');* Where T is the table containing the data, 'Group' represents the output variable, 'AV' represents average firing rate, 'Peak' represents peak firing rate, 'Bits' represents bits per second, 'SI' represents spatial information content, 'SC' represents spatial coherence, 'size' represents place field size, 'Outratio' represents outfield/infield ratio, and 'Infield' represents infield firing rate. The input ‘binomial’ in the function indicates that the variable dependent is binary. The input ‘logit’ indicates that the function logit link was used to model the probability of the binary outcome. Predicted probabilities were extracted using the function *predict*. Later, they were used to generate a Receiver Operating Characteristic (ROC) curve and calculate the Area Under the Curve (AUC) to evaluate its discriminative ability using the function *perfcurve*. The AUC value measures the model's performance to distinguish between the presence and absence of renewal. Coefficient estimates and their p-values were extracted to identify significant predictors with a threshold of $p<0.05$.

**GLM for Context prediction:** Two logistic regression models were developed to classify contextual categories (context A vs. context B) using binned spike counts from PCs. Models were implemented using an adapted logistic regression function in MATLAB from the public GitHub repository “Machine Learning Algorithms from Scratch” by Dr. Milan Parmar (https://github.com/upul/Machine-Learning-Algorithms-From-Scratch). The target variable was the context category (A = 0, B = 1), with normalized spike counts (min-max normalisation) as predictors. Both models employed logistic regression with Iterative Reweighted Least Squares (IRLS), which iteratively minimizes prediction errors (Burrus, 2012). Training datasets comprised 70% of spike counts from day 1 acquisition (context A) and extinction sessions (context B), specifically chosen to minimize learning-related biases. The first model was tested using the remaining 30% of acquisition and extinction spike counts. The second model tested renewal spike counts (context A) against the remaining 30% of extinction data. Spike counts were randomly sampled, and models were independently applied 1000 times per group (Non-Renewers and Renewers). Model performance was evaluated based on accuracy, precision, recall, and AUC-ROC curves.

**Field Displacement** for each neuron between trials were calculated by first selecting the bin containing the peak of the field for each trial and neuron. Later, the differences between the bins containing the peaks of the desired trials were calculated to compare shifts between sessions. These displacement values were subsequently used to categorize the shifts of place fields in displacements of less than 10 bins or greater than 10 bins in any possible direction (forward or backward displacement in the maze).

**Spatial Correlations** between trials for each neuron were calculated to explore the degree of remapping between trials. The firing maps vectors were correlated using Pearson's correlation coefficient and the function “xcorr” in MATLAB. Correlation values ranged between -1 (max negative correlation) to 1 (max positive correlation).

**Statistical Analysis:** it was performed using MATLAB, with all statistical significances set at $p<0.05$. Normality was tested using the Kolmogorov-Smirnov test. To evaluate significance levels between sessions and behavioral groups, one-way or two-way analysis of variance with repeated measures (rmANOVA) was applied. Proportions of behavioral response types were compared using a chi-square test. For non-normally distributed variables, the Wilcoxon rank-sum test was utilized, while the two-sample Kolmogorov-Smirnov test was employed to compare distributions. For circular data (theta phase modulation of spikes), the Watson-Williams multi-sample test from the CircStat toolbox in MATLAB (11) was used. All tests were followed by post hoc Bonferroni comparisons or Wilcoxon signed-rank tests for pairwise comparisons with Bonferroni corrections. When Bonferroni corrections were applied, statistical significance was adjusted according to the number of comparisons. Cohen's d was calculated to measure effect sizes. Plots were generated in MATLAB using the gramm toolbox (12), with error bars representing mean ± 95% confidence intervals (computed via t-distribution).

**Histology:** Electrode placements were histologically confirmed with Nissl staining of coronal slices (30μm thickness) (13).

### SI References


1. F. Draht, *et al.*, Experience-Dependency of Reliance on Local Visual and Idiothetic Cues for Spatial Representations Created in the Absence of Distal Information. *Front. Behav. Neurosci.* **11** (2017).

2. G. Paxinos, C. Watson, *The Rat Brain in Stereotaxic Coordinates* (Elsevier Academic Press, 2005).

3. M. Méndez-Couz, J. M. Becker, D. Manahan-Vaughan, Functional Compartmentalization of the Contribution of Hippocampal Subfields to Context-Dependent Extinction Learning. *Front. Behav. Neurosci.* **13** (2019).

4. M. A. A. van der Meer, A. D. Redish, Low and High Gamma Oscillations in Rat Ventral Striatum have Distinct Relationships to Behavior, Reward, and Spiking Activity on a Learned Spatial Decision Task. *Front. Integr. Neurosci.* **3**, 9 (2009).

5. M. Valero, A. Navas-Olive, L. M. De La Prida, G. Buzsáki, Inhibitory conductance controls place field dynamics in the hippocampus. *Cell Rep.* **40**, 111232 (2022).

6. L. Dolón Vera, B. Dietz, D. Manahan-Vaughan, Distal but not local auditory information supports spatial representations by place cells. *Cereb. Cortex* **34**, bhae202 (2024).

7. É. Duvelle, *et al.*, Insensitivity of Place Cells to the Value of Spatial Goals in a Two-Choice Flexible Navigation Task. *J. Neurosci.* **39**, 2522–2541 (2019).

8. R. M. Grieves, E. R. Wood, P. A. Dudchenko, Place cells on a maze encode routes rather than destinations. *eLife* **5**, e15986 (2016).

9. R. E. Harvey, *et al.*, Linear Self-Motion Cues Support the Spatial Distribution and Stability of Hippocampal Place Cells. *Curr. Biol.* **28**, 1803-1810.e5 (2018).

10. A. Fernández-Ruiz, *et al.*, Entorhinal-CA3 Dual-Input Control of Spike Timing in the Hippocampus by Theta-Gamma Coupling. *Neuron* **93**, 1213-1226.e5 (2017).

11. P. Berens, **CircStat** : A *MATLAB* Toolbox for Circular Statistics. *J. Stat. Softw.* **31** (2009).

12. P. Morel, Gramm: grammar of graphics plotting in Matlab. *J. Open Source Softw.* **3**, 568 (2018).

13. D. Manahan-Vaughan, Priming of group 2 metabotropic glutamate receptors facilitates induction of long-term depression in the dentate gyrus of freely moving rats. *Neuropharmacology* **37**, 1459–1464 (1998).

**Figure S1**.

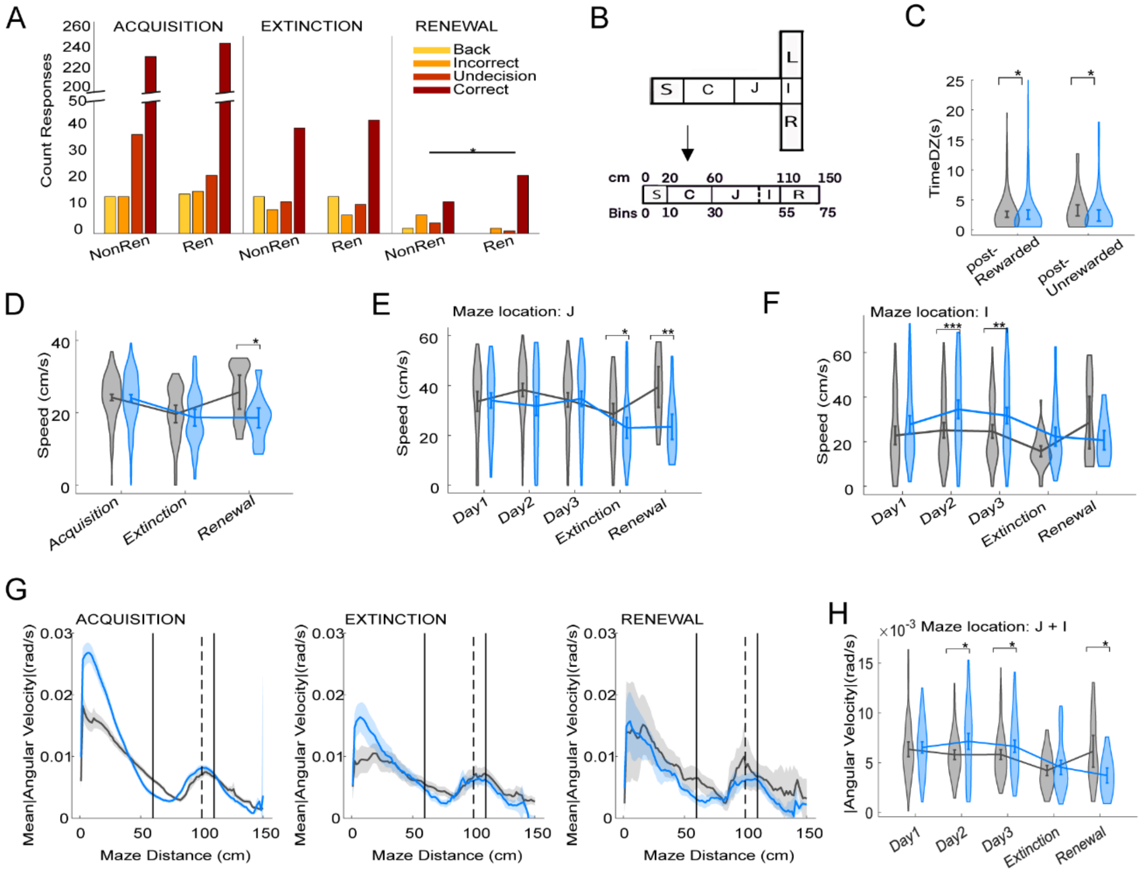


**Figure S1**. **Nuances of behavioural patterns by group.**

(**A)** Count of trial types (Back, Incorrect, Indecision, Correct -see Methods for trial classification) across phases. Data in the x-axis are grouped into Non-Renewers (NonRen) and Renewers (Ren) groups.

**(B)** Representation of maze areas and linearization strategy. Top: division of maze segments with S -Starting-, C -Corridor-, J -Junction-, I -Intersection-, L – left Arm or R-right Arm. **Bottom**: linearization of data represented in cm scale and in bins (2 cm/bin).

(**C**) Violin plots of time spent in the decision zone (i.e. J and I segments of T-maze) during trials where the correct arm was chosen during acquisition, but where the choice was followed by rewarded (post-Rewarded) or unrewarded (post-Unrewarded) outcomes (Data from Day 2 and Day 3).

**(D–F)** Violin plots of average speed **(D)**, speed in junction **(E)** and intersection **(F)**.

**(G**) Line plots of angular velocity across the maze and phase.

(**H**) Violin plots of angular velocity in DZ (J + I). Error bars represent mean ± 95% confidence interval (CI). Asterisks indicate significance (*, p<0.05). See SI Appendix Table S1 for statistical analyses.

**Figure S2.**

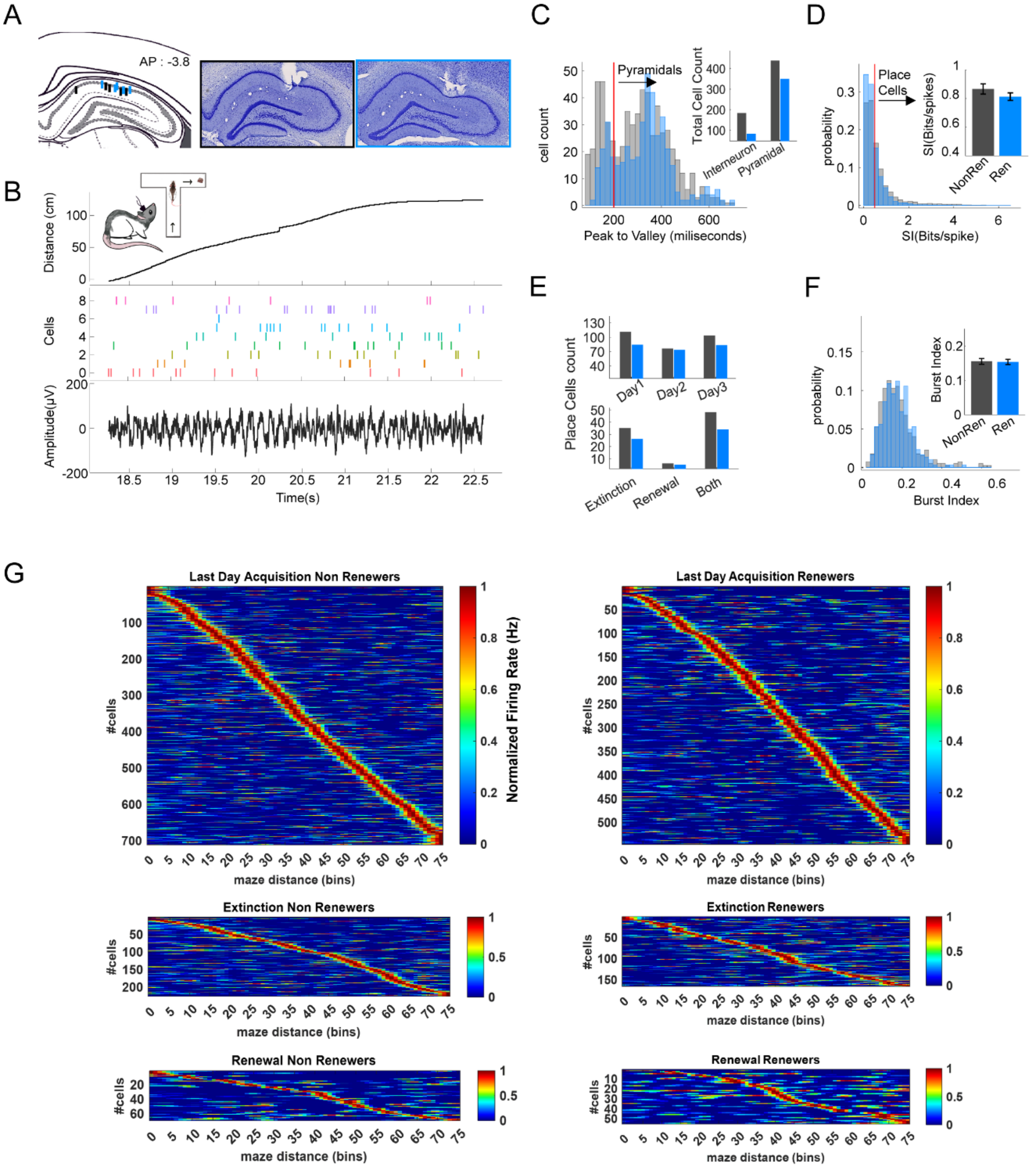


**Fig. S2. Place cell classification, controls, and population metrics.**

(**A**) Left: coronal atlas section showing all electrode placements. **Left**: example of Nissl-stained sections with tetrode tracks in CA1 (black, non-renewers; blue, renewers).(
**B**) Example of trial showing linearized position, spiking activity of 8 place cells, and corresponding local field potentials (LFPs).

(**C**) **Left**: histogram of spike peak-to-valley durations, with interneuron threshold (red dashed line). **Right**: counts of pyramidal cells vs interneurons per group (no group difference: $\chi^2(2, N=1052)=2$, $p=0.157$).

(**D**) Distribution of spatial information scores used for place cell classification (threshold = 0.5 bits/spike, red dashed line) with inset showing similar means across groups ($Z=-1.636$, $p=0.107$).

(**E**) top: counts of place cells by acquisition day, showing stable numbers across days (**day 1**: NonRen=104, Ren=73; **day 2**: NonRen=73, Ren=67; **day 3:** NonRen=89, Ren=80). **Bottom**: counts of place cells that were active only during EL, only during renewal, or during both events (**day 4**: NonRen=79 total, with 31 only EL, 5 only renewal, 43 both; R=61 total, with 23 only EL, 5 only renewal, 33 both; no differences: $\chi^2(2, N=154)=6$, $p=0.199$).

(**F**) Burst index distributions by group (Non-Renewers: 0.155 [0.147–0.163]; Renewers: 0.154 [0.146–0.161]; $Z=-1.013$, $p=0.311$), indicating similar discharge dynamics.

(**G**) Aligned, normalized firing rate maps for all PCs on day 3 (top), EL (middle), and renewal (bottom), sorted by peak location, for non-renewers (left) and renewers (right).

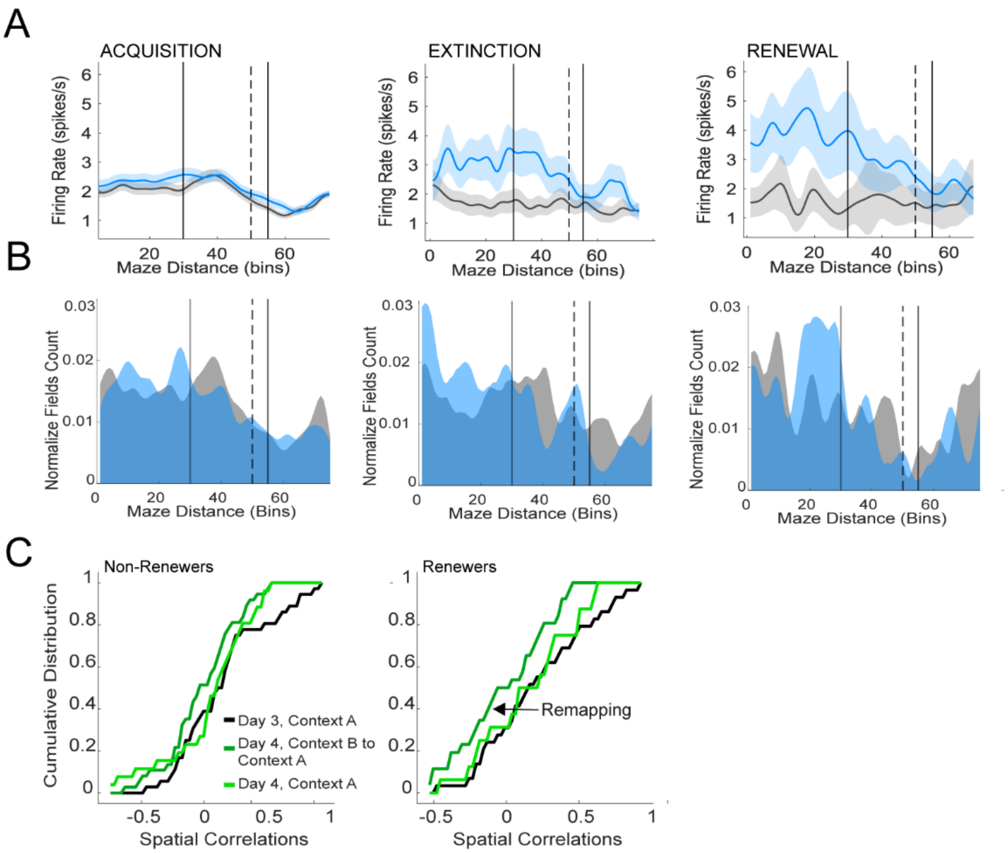


**Figure S3. Additional features of place cell population dynamics.**

(**A**) Mean firing rates and (**B**) Place field counts by maze bin across phases for Renewers (blue) and Non-Renewers (grey). Shadow error in (A) represents confidence interval (CI).

Note the sharp remapping in Renewers during extinction learning and renewal, contrasting with stable distributions in Non-Renewers.

(**C**) Cumulative distributions of spatial correlations of firing maps showed in Fig. S3B. Note the decrease on spatial correlations for Day 4, Context B to Context A in Renewers, distribution that was lower and significantly different from Day 4, Context A (renewal) and Day 3, Context A (acquisition), indicating a remapping of place fields during the context change in day 4. See SI Appendix **Table S7** for statistical analyses.

**Table for Figure 1**

| Figure Panel | Variable | Group/comparison | Mean ± SEM | Test | Test stat | p-value | Notes |
|---|---|---|---|---|---|---|---|
| 1B | **Correct responses over sessions** | All animals | | rmANOVA | F(15,135) = 7.034 | p < 0.001 | Significant session effect |
| | **Correct responses Session 1 (Acquisition)** | All animals | 3.2 ± 0.51 | | | | Baseline acquisition |
| | **Correct responses Session 12 (Acquisition)** | All animals | 4.8 ± 0.13 | Post hoc Bonferroni, S1 * S12 | | p < 0.05 | Indicates learning |
| | **Correct responses Session 13 (EL)** | All animals | 3.4 ± 0.33 | | | | Generalization to new context B |
| | **Correct responses Session 15 (EL)** | All animals | 1.5 ± 0.22 | Post hoc Bonferroni, S12 * S15 | | p < 0.001 | Indicates extinction |
| 1C | **Renewal classification** | Renewers: N=5, Non-Renewers: N=5 | | ≥80% increase from S15 to S16 | | | 50% animals exhibited renewal |
| | **Correct responses over sessions by groups** | | | rmANOVA, Session*Group | F(1,15) = 1.231 | p = 0.258 | No interaction across sessions & group |
| | | | | Post hoc Bonferroni, S1-S15 | | p > 0.05 | No group differences in acquisition or EL |
| | **Correct responses * S16* Group** | Renewers | 4.40 ± 0.40 | Post hoc Bonferroni, S16*Group | | p < 0.05 | Significant difference at Session 16 |
| | | Non-Renewers | 2.40 ± 0.51 | | | | |

| | | Group/comparison | | Test | Statistic | p-value | Effect size (d) |
|---|---|---|---|---|---|---|---|
| 1D | **Latency** | Non-Renewers Mean 95%[CI]) | Renewers (Mean 95%[CI]) | | | | |
| | Acquisition | 6.82 (5.84 – 7.80) | 5.96(4.89 – 7.02) | Wilcoxon rank-sum test | Z = 2.074 | p < 0.05 | d = 0.108 |
| | EL | 7.49(4.21 – 10.78) | 7.47(5.19 – 9.75) | Wilcoxon rank-sum test | Z = -1.315 | p = 0.188 | d = 0.002 |
| | Renewal | 7.33(2.21 – 12.45) | 10.0 (5.44 – 14.48) | Wilcoxon rank-sum test | Z = -0.721 | p = 0.471 | d=-0.276 |
| | Non-Renewers: Acquisition * EL | | | | Z = 0.528 | p = 0.059 | d = -0.086 |
| | Non-Renewers: Acquisition * Renewal | | | | Z = -0.194 | p = 0.846 | d = -0.068 |
| | Non-Renewers: EL*Renewal | | | | Z = -0.522 | p = 0.601 | d = 0.017 |
| | Renewers: Acquisition * EL | | | | Z = -2.531 | p = 0.01 | d = -0.184 |
| | Renewers: Acquisition * Renewal | | | | Z = -9.964 | p < 0.01 | d = -0.469 |
| | Renewers: EL*Renewal | | | | Z = -0.769 | p = 0.441 | d = -0.294 |
| 1E | **Time in decision zone** | | | | | | |
| | Acquisition | 2.94(2.48 - 3.41) | 2.71(2.22 - 3.21) | Wilcoxon rank-sum test | Z = 3.111 | p < 0.01 | d = 0.062 |
| | EL | 5.46(3.15 – 7.72) | 4.92(3.26 – 6.57) | Wilcoxon rank-sum test | Z = 0.686 | p = 0.493 | d = 0.087 |
| | Renewal | 2.89(1.38 – 4.40) | 7.4(0.94 - 13.85) | Wilcoxon rank-sum test | Z = -1.099 | p = 0.271 | d = -0.378 |
| | Non-Renewers: Acquisition * EL | | | Wilcoxon rank-sum test | Z = -3.969 | p < 0.001 | d = -0.597 |
| | Non-Renewers: Acquisition * Renewal | | | Wilcoxon rank-sum test | Z = -0.571 | p = 0.568 | d = 0.014 |
| | Non-Renewers: EL*Renewal | | | Wilcoxon rank-sum test | Z = 1.681 | p = 0.092 | d = 0.408 |
| | Renewers: Acquisition * EL | | | Wilcoxon rank-sum test | Z = -3.857 | p < 0.001 | d = -0.531 |
| | Renewers: Acquisition * Renewal | | | Wilcoxon rank-sum test | Z = -3.856 | p < 0.001 | d = -0.846 |
| | Renewers: EL*Renewal | | | Wilcoxon rank-sum test | Z = -0.256 | p = 0.797 | d = -0.261 |
| 1F | **Time DZ session 12** | 2.37(1.51 – 3.22) | 1.32(0.96 – 1.67) | | | | |
| | **Time DZ session 13** | 3.25(1.97 – 4.54) | 5.50(2.27 – 8.73) | | | | |
| | **Time DZ session 8** | 3.01(1.13 - 4.88) | 2.66(1.05 - 4.28) | | | | |
| | **Time DZ session 9** | 3.19(1.75 - 4.63) | 1.87(1.13 - 2.57) | | | | |
| | **Non- Renewers: Session 12 VS Session 13** | | | Wilcoxon rank-sum test | Z = -1.423 | p = 0.154 | d = -0.423 |
| | **Renewers: Session 12 VS Session 13** | | | Wilcoxon rank-sum test | Z = -2.733 | p < 0.01 | d = -0.965 |
| | **Non- Renewers: Session 8 VS Session 9** | | | Wilcoxon rank-sum test | Z = -0.088 | p = 0.929 | d = -0.049 |
| | **Renewers: Session 8 VS Session 9** | | | Wilcoxon rank-sum test | Z = 1.145 | p = 0.252 | d = 0.363 |
| 1G | **Bin Location of minimun speed (in DZ)** | | | | | | |
| | Acquisition | 77.04(71.84 - 82.23) | 74.82(70.27 - 79.38) | Wilcoxon rank-sum test | Z = 3.244 | p < 0.001 | d = 0.058 |
| | EL | 90.29(80.95 - 99.63) | 80.74(72.23 - 89.26) | Wilcoxon rank-sum test | Z = 3.576 | p < 0.001 | d = 0.346 |
| | Renewal | 93.52(80.56 - 106.48) | 90.77(85.20 - 96.34) | Wilcoxon rank-sum test | Z = 1.265 | p = 0.205 | d = 0.173 |
| 1H | **Bin Location of Angular velocity (in DZ)** | | | | | | |
| | **Bin of angular velocity increase onset, Acquisition** | 86.21(84.76 - 87.66) | 74.16(72.84 - 75.49) | Wilcoxon rank-sum test | Z= 12.556 | p < 0.001 | d = 1.121 |
| | **Bin of maximum angular velocity, Acquisition** | 91.99(89.722 - 94.23) | 98.23(97.03 - 99.53) | Wilcoxon rank-sum test | Z = -2.59 | p < 0.01 | d = -0.454 |

**Table S1. Statistics Overview for Figure 1**

**Table for Figure S1**

| Figure Panel | Variable | Group/comparison | Test | Test stat | p-value |
|---|---|---|---|---|---|
| S1A | | | | | |
| | **Alternative response proportions (Acquisition)** | Renewers vs Non-Renewers | Chi-square | $X^2(2, N=600)= 4.932$ | $p = 0.177$ |
| | **Alternative response proportions (EL)** | Renewers vs Non-Renewers | Chi-square | $X^2(2, N=150)= 0.402$ | $p = 0.923$ |
| | **Alternative response proportions (Renewal)** | Renewers vs Non-Renewers | Chi-square | $X^2(2, N=50)= 9.519$ | $p < 0.05$ |

| Figure Panel | Variable | Group/comparison | | Test | Statistic | p-value | Effect size (d) |
|---|---|---|---|---|---|---|---|
| | | Non-Renewers Mean 95%[CI]) | Renewers (Mean 95%[CI]) | | | | |
| S1C | | | | | | | |
| | **Time in DZ after correct-non-rewarded trial (post-Unrewarded)** | 3.24(2.33 – 4.16) | 2.39(1.47 – 3.29) | Wilcoxon rank-sum test | Z = 2.556 | p < 0.05 | d = 0.271 |
| | **Time in DZ after correct-rewarded trial (post-Rewarded)** | 2.58(2.07 – 3.08) | 2.70(1.89 – 3.51) | Wilcoxon rank-sum test | Z = 2.201 | p < 0.05 | d = -0.033 |
| | **Proportion of alternative responses after non-rewarded trial** | | | Chi-square | $X^2(2, N=109) = 3.596$ | p = 0.309 | |
| S1D | **Speed Average** | | | | | | |
| | Acquisition | 23.36(22.35 – 24.36) | 23.18(22.14 – 24.22) | Wilcoxon rank-sum test | Z = 0.196 | p = 0.845 | d = 0.022 |
| | EL | 19.26(16.76 – 21.76) | 18.67( 16.37 – 20.96) | Wilcoxon rank-sum test | Z = 0.797 | p = 0.425 | d = 0.078 |
| | Renewal | 25.70(21.01 – 30.40) | 18.60(15.87 – 21.33) | Wilcoxon rank-sum test | Z = 2.541 | p < 0.05 | d = 1.074 |
| | Non-Renewers: Acquisition * EL | | | Wilcoxon rank-sum test | Z = 3.330 | p <0.001 | d = 0.523 |
| | Non-Renewers: Acquisition * Renewal | | | Wilcoxon rank-sum test | Z = -0.921 | p = 0.356 | d = -0.299 |
| | Non-Renewers: EL*Renewal | | | Wilcoxon rank-sum test | Z = -2.204 | p < 0.05 | d = -0.834 |
| | Renewers: Acquisition * EL | | | Wilcoxon rank-sum test | Z = 3.862 | p < 0.001 | d = 0.552 |
| | Renewers: Acquisition * Renewal | | | Wilcoxon rank-sum test | Z = 3.280 | p < 0.001 | d = 0.562 |
| | Renewers: EL*Renewal | | | Wilcoxon rank-sum test | Z = 0.076 | p = 0.939 | d = 0.009 |
| S1E | **Speed Average Junction Zone** | | | | | | |
| | Acquisition | 37.26(35.72 - 38.81) | 34.95(33.19 - 36.72) | Wilcoxon rank-sum test | Z= 1.379 | p = 0.167 | d = 0.179 |
| | EL | 29.95(25.91 - 33.99) | 23.05(18.95 - 27.16) | Wilcoxon rank-sum test | Z= 2.528 | p < 0.01 | d = 0.539 |
| | Renewal | 39.47(31.30 - 47.65) | 23.45(18.40 - 28.49) | Wilcoxon rank-sum test | Z = 3.261 | p < 0.001 | d = 1.345 |
| | Non-Renewers: Acquisition * EL | | | Wilcoxon rank-sum test | Z = 3.381 | p < 0.001 | d = 0.623 |
| | Non-Renewers: Acquisition * Renewal | | | Wilcoxon rank-sum test | Z = -0.488 | p = 0.625 | d = -0.188 |
| | Non-Renewers: EL*Renewal | | | Wilcoxon rank-sum test | Z = -2.081 | p < 0.05 | d = -0.774 |
| | Renewers: Acquisition * EL | | | Wilcoxon rank-sum test | Z = 4.813 | p < 0.001 | d = 0.865 |
| | Renewers: Acquisition * Renewal | | | Wilcoxon rank-sum test | Z = 3.619 | p < 0.001 | d = 0.843 |
| | Renewers: EL*Renewal | | | Wilcoxon rank-sum test | Z = -0.173 | p = 0.862 | d = -0.031 |
| S1F | **Speed Average Intersection Zone** | | | | | | |
| | Acquisition | 25.60(23.68 - 27.52) | 32.85(30.66 - 35.04) | Wilcoxon rank-sum test | Z = -4.863 | p < 0.001 | d = -0.455 |
| | EL | 16.78(14.57 - 18.99) | 22.27(18.14 - 26.41) | Wilcoxon rank-sum test | z = -1.650 | P = 0.098 | d = -0.506 |
| | Renewal | 28.59(16.89 - 40.28) | 20.67(16.37 - 24.97) | Wilcoxon rank-sum test | Z = 0.738 | p = 0.46 | d = 0.593 |
| | Non-Renewers: Acquisition * EL | | | Wilcoxon rank-sum test | Z = 3.220 | p < 0.001 | d = 0.644 |
| | Non-Renewers: Acquisition * Renewal | | | Wilcoxon rank-sum test | Z = -0.317 | p = 0.751 | d = -0.202 |
| | Non-Renewers: EL*Renewal | | | Wilcoxon rank-sum test | Z = -1.499 | p = 0.133 | d = -1.113 |
| | Renewers: Acquisition * EL | | | Wilcoxon rank-sum test | Z = 3.948 | p < 0.001 | d = 0.637 |
| | Renewers: Acquisition * Renewal | | | Wilcoxon rank-sum test | Z = 3.276 | p < 0.001 | d = 0.732 |
| | Renewers: EL*Renewal | | | Wilcoxon rank-sum test | Z = 0.076 | p = 0.939 | d = 0.130 |
| S1H | **Average Angular velocity in DZ** | | | | | | |
| | **Day 2** | .006(.005-.006) | .007(.006-.008) | Wilcoxon rank-sum test | Z = -2.733 | p < 0.01 | d = -0.469 |
| | **Day 3** | .006(.005 - .006) | .007(.006 - .007) | Wilcoxon rank-sum test | Z = -2.031 | p < 0.05 | d = -0.323 |
| | **Renewal** | .007(.005 - .009) | .004(.003-.005) | Wilcoxon rank-sum test | Z = 2.396 | p < 0.05 | d = 1.209 |

### Table S2. Statistics Overview for Figure S1

PLACE CELLS SPATIAL PROPERTIES

| | | Day 1 | Day 2 | Day 3 | Day 4 Exctinction | Day 4 Renewal |
|---|---|---|---|---|---|---|
| Peak firing rate (spikes/s) | Non-Renewers | 9.34 (8.84 - 9.84) | 8.94 (8.44 - 9.44) | 8.58 (8.17 - 9.00) | 6.94 (6.37 - 7.50) | 7.75 (6.37 - 9.13) |
| | Renewers | 9.44 (8.82 - 10.07) | 9.14 (8.65 - 9.62) | 10.23 (9.72 - 10.74) | 11.88 (10.92 - 12.83) | 9.45 (8.12 - 10.77) |
| Average Firing Rate | Non-Renewers | 1.61 (1.51 - 1.71) | 1.74 (1.63 - 1.85) | 1.74 (1.66 - 1.82) | 1.54 (1.41 - 1.67) | 1.54 (1.26 - 1.82) |
| | Renewers | 1.73 (1.61 - 1.85) | 1.89 (1.78 - 2.00) | 1.9 (1.80 - 1.99) | 2.37 (2.17 - 2.57) | 2.23 (1.94 - 2.51) |
| Spatial Information (Bits/spike) | Non-Renewers | 1.17 (1.09 - 1.24) | 0.84 (0.77 - 0.90) | 0.79 (0.75 - 0.84) | 0.76 (0.67 - 0.85) | 0.78 (0.66 - 0.91) |
| | Renewers | 0.76 (0.71 - 0.81) | 0.85 (0.80 - 0.90) | 0.91 (0.86 - 0.96) | 0.99 (0.90 - 1.08) | 0.64 (0.56 - 0.72) |
| Spatial Coherence | Non-Renewers | 2.86 (2.82 2.90) | 2.73 (2.69 - 2.78) | 2.73 (2.69 - 2.76) | 2.65 (2.59 - 2.71) | 2.72 (2.62 - 2.83) |
| | Renewers | 2.82 (2.77 2.87) | 2.79 (2.75 - 2.83) | 2.82 (2.79 - 2.85) | 2.88 (2.82 - 2.95) | 2.68 (2.57 - 2.79) |
| Field Width (Bins) | Non-Renewers | 11.47 (10.93 - 12.02) | 10.9 (10.25 - 11.55) | 11.05 (10.56 - 11.54) | 11.85 (10.88 - 12.82) | 9.47 (8.33 - 10.61) |
| | Renewers | 10.84 (10.19 - 11.50) | 11.43 (10.83 - 12.02) | 11.03 (10.53 - 11.52) | 12.12 (11.08 - 13.17) | 14.78 (12.46 - 17.10) |
| Infield Firing Rate | Non-Renewers | 6 (5.68 - 6.32) | 5.66 (5.35 - 5.97) | 5.44 (5.17 - 5.71) | 4.37 (3.99 - 4.75) | 4.86 (3.98 - 5.73) |
| | Renewers | 6.01 (5.62 - 6.40) | 5.82 (5.52 - 6.13) | 6.52 (6.19 6.85) | 7.63 (7.00 - 8.27) | 5.88 (5.06 - 6.71) |
| Out/Infield Ratio | Non-Renewers | 0.15 (0.14 - 0.16) | 0.17 (0.16 - 0.19) | 0.19 (0.18 - 0.20) | 0.26 (0.23 - 0.28) | 0.22 (0.18 - 0.27) |
| | Renewers | 0.2 (0.19 - 0.22) | 0.21 (0.19 - 0.22) | 0.19 (0.18 - 0.20) | 0.23 (0.21 - 0.25) | 0.29 (0.26 - 0.33) |
| Bits/second | Non-Renewers | 1.46 (1.36 - 1.56) | 1.15 (1.08 - 1.22) | 1.21 (1.14 - 1.27) | 0.95 (0.85 - 1.06) | 1.01 (0.83 - 1.19) |
| | Renewers | 1.18 (1.10 - 1.27) | 1.4 (1.32 - 1.47) | 1.48 (1.40 - 1.56) | 2.17 (1.96 - 2.38) | 1.45 (1.19 - 1.70) |
| Bin Location | Non-Renewers | 31.91 (30.17 - 33.64) | 33.74 (31.58 - 35.89) | 34.52 (32.73 - 36.31) | 34.27 (31.39 - 37.15) | 33.01 (26.59 - 39.43) |
| | Renewers | 29 (26.70 - 31.30) | 33.19 (31.10 - 35.29) | 33.16 (31.44 - 34.88) | 28.62 (25.49 - 31.75) | 28.99 (23.51 - 34.47) |

**Table S3. Summary of place field properties.**

All values are presented as mean ± 95% confidence interval (CI).

Table for Figure 2

| Figure Panel | Variable | Group/comparison<br>Non-Renewers Mean 95%[CI]) | <br>Renewers (Mean 95%[CI]) | Test | Statistic | p-value | Effect size (d) |
|---|---|---|---|---|---|---|---|
| **2B** | **Peak FR** | | | | | | |
| | Acquisition | 9.4 (9.1–10.0) | 9.95(9.62 - 10.27) | Wilcoxon rank-sum test | Z = -2.323 | p < 0.05 | d = -0.069 |
| | EL | 7.2 (6.6–7.8) | 12.21(11.24 - 13.18) | Wilcoxon rank-sum test | Z = -8.189 | p < 0.001 | d = -1.011 |
| | Renewal | 7.75(6.37 - 9.13) | 10.51(8.97 - 12.05) | Wilcoxon rank-sum test | Z = -2.591 | p < 0.01 | d = -0.512 |
| | Non-Renewers: Acquisition * EL | | | Wilcoxon rank-sum test | Z = 6.032 | p < 0.001 | d = 0.419 |
| | Non-Renewers: Acquisition * Renewal | | | Wilcoxon rank-sum test | Z = 2.431 | p < 0.01 | d = -0.308 |
| | Non-Renewers: EL*Renewal | | | Wilcoxon rank-sum test | Z = -0.579 | p = 0.563 | d = -0.134 |
| | Renewers: Acquisition * EL | | | Wilcoxon rank-sum test | Z = -4.475 | p < 0.001 | d = -0.404 |
| | Renewers: Acquisition * Renewal | | | Wilcoxon rank-sum test | Z = -0.628 | p = 0.592 | d = -0.1 |
| | Renewers: EL*Renewal | | | Wilcoxon rank-sum test | Z = 1.971 | p = 0.048 | d = 0.290 |
| **2C** | **Bits/Second** | | | | | | |
| | Acquisition | 1.39(1.34 - 1.45) | 1.41(1.36 - 1.45) | Wilcoxon rank-sum test | Z = -3.085 | p < 0.05 | d = -0.011 |
| | EL | 1(0.88 - 1.12) | 2.25(2.03 - 2.47) | Wilcoxon rank-sum test | Z = -10.196 | p < 0.001 | d = -1.159 |
| | Renewal | 1.01(0.83 - 1.18) | 1.58(1.32 - 1.85) | Wilcoxon rank-sum test | Z = -3.161 | p < 0.01 | d = -0.681 |
| | Non-Renewers: Acquisition * EL | | | Wilcoxon rank-sum test | Z = 6.418 | p < 0.001 | d = 0.381 |
| | Non-Renewers: Acquisition * Renewal | | | Wilcoxon rank-sum test | Z = 2.914 | p < 0.05 | d = 0.371 |
| | Non-Renewers: EL*Renewal | | | Wilcoxon rank-sum test | Z = -0.658 | p = 0.512 | d = -0.008 |
| | Renewers: Acquisition * EL | | | Wilcoxon rank-sum test | Z = -7.905 | p < 0.001 | d = -0.951 |
| | Renewers: Acquisition * Renewal | | | Wilcoxon rank-sum test | Z = -0.986 | p = 0.324 | d = -0.219 |
| | Renewers: EL*Renewal | | | Wilcoxon rank-sum test | Z = 3.371 | p < 0.001 | d = 0.533 |
| **2D** | **SI (Bits/spike)** | | | | | | |
| | Acquisition | 0.92(0.88 - 0.95) | 0.84(0.81 – 0.87) | Wilcoxon rank-sum test | Z = 0.085 | p = 0.931 | d = 0.122 |
| | Day 3 | 0.77(0.73 - 0.81) | 0.9(0.85 - 0.94) | Wilcoxon rank-sum test | Z = -4.029 | p < 0.001 | d = -0.251 |
| | EL | 0.75(0.67 - 0.84) | 0.98(0.89 - 1.11) | Wilcoxon rank-sum test | Z = -5.193 | p < 0.001 | d = -0.391 |
| | Renewal | 0.78(0.66 - 0.91) | 0.63(0.56 - 0.71) | Wilcoxon rank-sum test | Z = 2.029 | p < 0.05 | d = 0.427 |
| | Non-Renewers: Acquisition * EL | | | Wilcoxon rank-sum test | Z = 3.939 | p < 0.001 | d = 0.233 |
| | Non-Renewers: Acquisition * Renewal | | | Wilcoxon rank-sum test | Z = 0.537 | p = 0.591 | d = 0.196 |
| | Non-Renewers: EL*Renewal | | | Wilcoxon rank-sum test | Z = -1.332 | p = 0.182 | d = -0.042 |
| | Non- Renewers: Day 1 * Day 3 | | Day 1: 1.14(1.06 - 1.22)<br>Day 3: 0.77(0.73 - 0.82) | Wilcoxon rank-sum test | Z = 6.746 | p < 0.001 | d = 0.528 |
| | Renewers: Acquisition * EL | | | Wilcoxon rank-sum test | Z = -3.554 | p<0.001 | d = -0.284 |
| | Renewers: Acquisition * Renewal | | | Wilcoxon rank-sum test | Z = 3.555 | p <0.000 | d = 0.438 |
| | Renewers: EL*Renewal | | | Wilcoxon rank-sum test | Z = 4.829 | p <0.001 | d = 0.744 |
| | Renewers: Day 1 * Day 3 | | Day 1: 0.74(0.69 - 0.79)<br>Day 3: 0.91(0.85 - 0.95) | Wilcoxon rank-sum test | Z = -4.252 | p < 0.001 | d = -0.321 |
| **2E** | **Spatial Coherence** | | | | | | |
| | Acquisition | 2.73(2.75 - 2.79) | 2.81(2.78 - 2.82) | Wilcoxon rank-sum test | Z = -1.976 | p = 0.048 | d = -0.107 |
| | EL | 2.65(2.59 - 2.71) | 2.88(2.81 - 2.95) | Wilcoxon rank-sum test | Z = -5.251 | p < 0.001 | d = -0.585 |
| | Renewal | 2.72(2.61 - 2.83) | 2.69(2.59 - 2.8) | Wilcoxon rank-sum test | Z = 0.722 | p = 0.470 | d = 0.067 |
| | Non-Renewers: Acquisition * EL | | | Wilcoxon rank-sum test | Z = 4.160 | p < 0.001 | d = 0.274 |
| | Non-Renewers: Acquisition * Renewal | | | Wilcoxon rank-sum test | Z = 0.901 | p = 0.363 | d = 0.109 |
| | Non-Renewers: EL*Renewal | | | Wilcoxon rank-sum test | Z = -1.372 | p = 0.184 | d = -0.177 |
| | Renewers: Acquisition * EL | | | Wilcoxon rank-sum test | Z = -2.221 | p < 0.05 | d = -0.212 |
| | Renewers: Acquisition * Renewal | | | Wilcoxon rank-sum test | Z = 2.287 | p < 0.05 | d = 0.288 |
| | Renewers: EL*Renewal | | | Wilcoxon rank-sum test | Z = 3.001 | p < 0.05 | d = 0.478 |
| **2F** | **Bin Width** | | | | | | |
| | Acquisition | 11.45(11.13 - 11.78) | 11.24(10.91 - 11.58) | Wilcoxon rank-sum test | Z = 0.028 | p = 0.98 | d = 0.035 |
| | EL | 12.06(11.06 - 13.07) | 12.33(11.26 - 13.38) | Wilcoxon rank-sum test | Z = -1.143 | p = 0.253 | d = -0.038 |
| | Renewal | 9.47(8.33 - 10.61) | 15.69(13.32 - 18.07) | Wilcoxon rank-sum test | Z = -3.934 | p < 0.001 | d = -0.894 |
| | Non-Renewers: Acquisition * EL | | | Wilcoxon rank-sum test | Z = -0.739 | p = 0.459 | d = -0.075 |
| | Non-Renewers: Acquisition * Renewal | | | Wilcoxon rank-sum test | Z = 2.37 | p = 0.017 | d = 0.337 |
| | Non-Renewers: EL*Renewal | | | Wilcoxon rank-sum test | Z = 2.46 | p = 0.013 | d = 0.416 |
| | Renewers: Acquisition * EL | | | Wilcoxon rank-sum test | Z = -2.382 | p < 0.05 | d = -0.228 |
| | Renewers: Acquisition * Renewal | | | Wilcoxon rank-sum test | Z = -4.074 | p < 0.01 | d = -0.854 |
| | Renewers: EL*Renewal | | | Wilcoxon rank-sum test | Z = -2.408 | p < 0.05 | d = -0.476 |
| **2G** | **Infield FR** | | | | | | |
| | Acquisition | 6.06(5.86 - 6.26) | 6.32(6.11 - 6.52) | Wilcoxon rank-sum test | Z = -2.562 | p < 0.01 | d = -0.071 |
| | EL | 4.51(4.12 - 4.91) | 7.79(7.16 - 8.42) | Wilcoxon rank-sum test | Z = -8.382 | p < 0.001 | d = -0.923 |
| | Renewal | 4.85(3.98 - 5.73) | 6.39(5.49 - 7.3) | Wilcoxon rank-sum test | Z = -2.598 | p < 0.01 | d = -0.471 |
| | Non-Renewers: Acquisition * EL | | | Wilcoxon rank-sum test | Z = 6.596 | p < 0.001 | d = 0.423 |
| | Non-Renewers: Acquisition * Renewal | | | Wilcoxon rank-sum test | Z = 2.749 | p < 0.01 | d = 0.328 |
| | Non-Renewers: EL*Renewal | | | Wilcoxon rank-sum test | Z = -0.547 | p = 0.584 | d = -0.119 |
| | Renewers: Acquisition * EL | | | Wilcoxon rank-sum test | Z = -4.438 | p<0.001 | d = -0.417 |
| | Renewers: Acquisition * Renewal | | | Wilcoxon rank-sum test | Z = -0.193 | p = 0.854 | d = -0.021 |
| | Renewers: EL*Renewal | | | Wilcoxon rank-sum test | Z = 2.466 | p < 0.05 | d = 0.377 |
| **2I** | **Theta power** | 0.05(0.04 - 0.06) | 0.07(0.06 - 0.07) | Wilcoxon rank-sum test | Z = -2.833 | p < 0.01 | d = -0.791 |
| | **Phase-lock** | | | Watson-Williams multi-sample test | F(1,226) = 3.26 | p = 0.072 | |
| | **Vector length** | | | Kruskal-Wallis One-way ANOVA | df = 2, $X^2$ = 13.39 | p<0.001 | |

### Table S4. Statistics Overview for Figure 2

| Variable | Group/comparison | | Test | Statistic | p-value | Effect size (d |
|---|---|---|---|---|---|---|
| | Non-Renewers Mean 95%[CI]) | Renewers (Mean 95%[CI]) | | | | |
| **Average FR** | | | | | | |
| Acquisition | 1.98(1.89 - 2.07) | 1.95(1.88 - 2.02) | Wilcoxon rank-sum test | Z = -2.377 | p < 0.05 | d = 0.022 |
| EL | 1.62(1.47 - 1.77) | 2.49(2.27 - 2.72) | Wilcoxon rank-sum test | Z = -6.521 | p < 0.001 | d = -0.727 |
| Renewal | 1.53(1.26 - 1.81) | 2.54(2.16 - 2.92) | Wilcoxon rank-sum test | Z = -4.061 | p < 0.001 | d = -0.803 |
| Non-Renewers: Acquisition * EL | | | Wilcoxon rank-sum test | Z = 2.488 | p < 0.05 | d = 0.214 |
| Non-Renewers: Acquisition * Renewal | | | Wilcoxon rank-sum test | Z = 1.827 | p = 0.067 | d = 0.258 |
| Non-Renewers: EL*Renewal | | | Wilcoxon rank-sum test | Z = 0.594 | p = 0.552 | d = 0.081 |
| Renewers: Acquisition * EL | | | Wilcoxon rank-sum test | Z = -4.897 | p < 0.001 | d = -0.439 |
| Renewers: Acquisition * Renewal | | | Wilcoxon rank-sum test | Z = -3.259 | p < 0.05 | d = -0.483 |
| Renewers: EL*Renewal | | | Wilcoxon rank-sum test | Z = -0.534 | p = 0.593 | d = -0.059 |
| **Outfield/Infield Ratios** | | | | | | |
| Acquisition | 0.17(0.17 - 0.18) | 0.20(0.19 - 0.21) | Wilcoxon rank-sum test | Z = -5.901 | p < 0.001 | d = -0.204 |
| EL | 0.25(0.23 - 0.27) | 0.23(0.21 - 0.25) | Wilcoxon rank-sum test | Z = 0.801 | p = 0.423 | d = 0.163 |
| Renewal | 0.22(0.17 - 0.27) | 0.29(0.26 - 0.33) | Wilcoxon rank-sum test | Z = -2.873 | p < 0.01 | d = -0.511 |
| Non-Renewers: Acquisition * EL | | | Wilcoxon rank-sum test | Z = -6.785 | p <0.001 | d = -0.616 |
| Non-Renewers: Acquisition * Renewal | | | Wilcoxon rank-sum test | Z= -1.985 | p = 0.046 | d = -0.383 |
| Non-Renewers: EL*Renewal | | | Wilcoxon rank-sum test | Z= 1.343 | p = 0.179 | d = 0.197 |
| Non- Renewers: Day 1 * Day 3 | | Day 1: 0.15(0.13 - 0.16)<br>Day 3: 0.19(0.18 - 0.21) | Wilcoxon rank-sum test | Z = -5.764 | p < 0.001 | d = -0.367 |
| Renewers: Acquisition * EL | | | Wilcoxon rank-sum test | Z = -3.037 | p < 0.01 | d = -0.244 |
| Renewers: Acquisition * Renewal | | | Wilcoxon rank-sum test | Z = -5.441 | p < 0.001 | d = -0.782 |
| Renewers: EL*Renewal | | | Wilcoxon rank-sum test | Z = -3.328 | p < 0.001 | d = -0.531 |
| Renewers: Day 1 * Day 3 | | Day 1: 0.02(0.19 - 0.22)<br>Day 3: 0.19(0.18 - 0.21) | Wilcoxon rank-sum test | Z = 1.601 | p = 0.109 | d = 0.125 |

**Table S5. Statistics  Overview for additional place field features.**

**Table for Figure 3**

| Figure panel | Comparison | Group | | Test | Statistic | p-value | Notes |
|---|---|---|---|---|---|---|---|
| 3A | **Acquisition (bins 1–75)** | Renewers vs. Non-Renewers | | Kolmogorov-Smirnov | K = 0.226 | p<0.01 | Spike distributions differed across most bins, except bins 8–18 and 70–75 |
| | **Acquisition vs. EL** | Non-Renewers | | Bin-wise comparisons | | all p < 0.05 | Overrepresentation reduced, spike activity increased in decision areas |
| | **Acquisition vs. EL** | Renewers | | Bin-wise comparisons | | all p < 0.05 | General redistribution, suggesting global remapping |
| 3B | **Bin location shift** | | | | | | |
| | **Day 3, Context A (first vs last trial)** | Renewers (cells = 29)<br>Non-Renewers (cells = 36) | | | | | Forward shift observed, not significant |
| | **Day 4, context B (first vs last trial)** | Renewers (cells = 43)<br>Non-Renewers (cells = 54) | | Wilcoxon rank-sum test | Renewers: Z = 1.905, d = 0.385 | p < 0.05 | Significant backward shift |
| | **Day 4, Context B to Context A (last vs first trial)** | Renewers (cells = 33)<br>Non-Renewers (cells = 46) | | Wilcoxon rank-sum test | Renewers: Z = -1.976, d = -0.48 | p < 0.05 | Significant forward shift vs EL |
| | **Day 4, Context A (first vs last trial)** | Renewers (cells = 16)<br>Non-Renewers (cells = 26) | | | | all p > 0.05 | No significant shift |
| 3C | **Proportions of large vs small displacements** | | | Chi-square | | all p > 0.05 | No group differences in proportion of displacement categories (<10 bins vs >10 bins) |
| 3F | **GLM for context decoding** | | | | | | |
| | | **Metric & Dataset** | **Group (Mean (95% CI))** | | | | |
| | **Model 1 (acquisition context A vs B)** | Accuracy (Training Acq+Ext) | Non-Renewers (0.803 (0.801–0.804))<br>Renewers (0.871 (0.869–0.873)) | Wilcoxon rank-sum test | Z = -37.901 | <0.001 | Renewers showed higher context separation in training |
| | | Accuracy (Test Acq+Ext) | Non-Renewers (0.656 (0.654–0.658))<br>Renewers (0.684 (0.681–0.687)) | Wilcoxon rank-sum test | Z = -15.107 | <0.001 | Renewers also showed higher context separation on test |
| | **Model 2 (renewal context A vs B)** | Accuracy (Test Ren+Ext) | Non-Renewers (0.582 (0.578–0.584))<br>Renewers (0.484 (0.481–0.486)) | Wilcoxon | Z = 33.937 | <0.001 | Renewers' model failed (below chance), indicating remapping during renewal |

*Logistic regression models were run with 1000 independent iterations using randomised spike count samples to predict context identity from spike distributions. Mean prediction accuracies and 95% CI are reported.

**Table S6. Statistics Overview for Figure 3**

**Table for Fig. S3**

| Figure Panel | Comparison | Measure | Non-Renewers | Renewers | Test | p-value | Effect size (d) | Notes |
|---|---|---|---|---|---|---|---|---|
| **S3A** | **Acquisition vs EL (Starting zone)** | Firing rate | | 2.677 (2.141–2.394) vs 2.845 (2.469–3.222) | Wilcoxon Z = -3.411 | <0.001 | -0.266 | |
| **S3A** | **Acquisition vs EL (Decision zone)** | Firing rate | | 2.201 (2.028–2.372) vs 3.025 (2.463–3.588) | Wilcoxon Z = -3.220 | <0.001 | -0.275 | |
| **S3A** | **Acquisition vs EL (Right arm)** | Firing rate | | 1.605 (1.525–1.688) vs 2.019 (1.794–2.289) | Wilcoxon Z = -2.849 | <0.01 | -0.287 | |
| **S3A** | **Decision zone Renewal vs Acquisition** | Firing rate | | 3.827 (3.020–4.633) vs 2.201 (2.028–2.372) | Wilcoxon Z = -5.071 | <0.001 | -0.552 | |
| **S3A** | **Decision zone Renewal vs EL** | Firing rate | | 3.827 (3.020–4.633) vs 3.025 (2.463–3.588) | Wilcoxon Z = -2.522 | <0.05 | -0.241 | |
| **S3B** | **Field distribution Acquisition** | | | | K = 0.120, p = 0.624 | 0.624 | | |
| **S3B** | **Field distribution EL** | K-S test Non-Renewers vs Renewers | | | K = 0.227, p = 0.035 | <0.05 | | |
| **S3B** | **Field distribution Renewal** | | | | K = 0.360, p < 0.001 | <0.001 | | |
| **S3C** | **Paired Spatial correlations** | | | | | | | |
| | **Renewers** | Day 4 Context B vs Day 3 Context A | | | KS = 0.400 | <0.001 | | Early decreased in EL |
| | **Renewers** | Day 4 Context B → A vs Day 3 Context A | | | KS = 0.340 | <0.01 | | Decreased during transition B → A |
| **S3C right** | **Renewers** | Day 4 Context A vs Day 3 Context A | | | KS = 0.220 | 0.077 | | Not significant |
| | **Non-Renewers** | Day 4 Context B vs Day 3 Context A | | | KS = 0.220 | 0.074 | | Not significant |
| | **Non-Renewers** | Day 4 Context B → A vs Day 3 Context A | | | KS = 0.260 | <0.05 | | Late decrease during transition B → A |
| **S3C left** | **Non-Renewers** | Day 4 Context A vs Day 3 Context A | | | KS = 0.240 | <0.05 | | Decrease in renewal |

**Table S7. Statistics Overview for Figure S3**
